# BenchIDPs: Evaluation of Conformational Preferences Molecular Mechanics and Solvent Interactions of IDPs across popular Force Fields and Water Models

**DOI:** 10.1101/2025.09.10.675321

**Authors:** Vishal Kumar, Mrudul Mete, Anupama Sharma, Prathit Chatterjee, U. Deva Priyakumar

**Author notes:** Contributing authors.

## Abstract

Intrinsically disordered proteins (IDPs), unlike globular proteins, lack stable secondary structure and exist as dynamic ensembles of conformations in physiological conditions. These conformations allow them to adapt to many roles while interacting with other proteins, forming partially folded soluble oligomers or insoluble plaques rich in ***β***-sheet, and are responsible for different pathological diseases. The dynamic nature of IDPs and the lack of well-defined binding sites make drug discovery and development both challenging. Although computational protocols, predominantly involving molecular dynamics (MD) simulations studies, have been complementing in understanding the structure and dynamics of IDPs, the choice of force fields become important in reproducing experimental observations. We herewith provide a systematic study to investigate the structural propensity of four IDPs (A***β***42, Tau43, amylin and ***α***S), using 13 combinations of force field-water models (FF-wm). In addition to validating with NMR observables, we also examine the water dynamics surrounding the chosen IDPs to underscore the generalizability of chosen FF-wms. C36-IDPSFF with amber99sb-dispersion water model (A99SB-disp) is found to perform optimal in simulating IDPs in comparison with other FF-wm combination based on structural propensity and comparison with experimental NMR **^3^**J***_HN−Hα_***-Coupling data as well as dynamical observables.

## 1 Introduction

Peptides and proteins are first synthesized as mostly unstructured amino acid chains that fold into distinct three-dimensional shapes with specific functions [1]. This process, known as protein folding, is essential for cellular function. The folding process involves overcoming kinetic entrapments from folded intermediate states, with decreased conformational heterogeneity as the folding process progresses [2]. Despite their tendency to form stable native structures, globular proteins may unfold under various conditions, such as high temperature, pH changes, mutations, or exposure to denaturants [3–6], making the misfolded or partially folded states energetically more favorable than their native counterparts. The aggregated states situated at relatively deeper energy minima in the non-native protein folding funnel often adopt cross-*β* sheet structures, characteristic of amyloidogenic states [7].

Misfolding and aggregation, commonly associated with amyloids, are involved in diseases such as Alzheimer’s, Parkinson’s, renal amyloidosis, and Type-II diabetes. Intrinsically disordered proteins, peptides, and regions (IDP/R) [8], because of their conformational flexibility arising from ruggedness in their energy landscape, have an inherent tendency to adopt intermolecular interactions resulting in (unfolded/partially folded) soluble and insoluble oligomers (amorphous or *β*-plaques rich aggregates). Although these have specific cellular functions such as signal transmission and vesicular trafficking in their soluble monomeric states, their oligomeric ensembles eventually have direct implications in cell-biology pathogenesis. Characterizing IDPs is particularly challenging due to their dynamic and unfolded nature. Conventional structure determination methods, such as X-ray crystallography and cryo-EM, can be difficult to apply to soluble IDPs [9]. Valuable insights can therefore be gained through spectroscopic techniques such as nuclear magnetic resonance (NMR) [10], circular dichroism (CD) spectroscopy [11], small-angle X-ray scattering (SAXS) [12], and Forster resonance energy transfer (FRET) [13]. However, additional atomistic details are needed to more accurately capture the conformational propensity [14]. This can be obtained from molecular dynamics (MD) simulations which provides bottoms-up approach to provide high-resolution in undertaking molecular level understanding of IDP conformational ensembles [15]. MD studies complement experiments to provide insights into the native and pathological functions. Considerable efforts have been made to accurately reproduce experimentally observed structural parameters (radius of gyration, secondary structure propensity [16, 17], etc.), on heterogeneous conformations involving IDPs. Additionally, the disordered nature of proteins, with minimal free energy differences between conformations, creates challenges in elucidating IDP phase-space [18]. Accurate predictions from MD trajectories require adequate sampling of these states. Consequently, long-timescale simulations are typically necessary [19], which in turn restricts the size of case-specific systems. Moreover, optimally large simulation boxes must be used to prevent self-interaction when employing periodic boundary conditions, which directly conflicts with the need for longer time-scale simulations. Despite thorough sampling efforts, limitations of conventional force fields or potential energy functions, can skew the molecular and thermodynamic properties between folded and unfolded states, potentially leading to an over-representation of either of the states.

Pertinent issues with the choice of force fields and water models include relatively high helical propensities compared to experimental observations [20, 21]. Also, weak protein-water interactions [22], in commonly used water models like TIP3P and TIP4P [23, 24], amplify inaccurate solvation-free energies [25],[26], and deviation from experimental observed conformations [27–29]. Therefore, accurate predictions lead to IDP observables with unconstrained atomistic simulations, remain a significant challenge [30]. This difficulty arises partly because current force fields were parameterized to describe the native states of folded (globular) proteins, with much less attention given to IDPs. Several studies have eventually underscored the limitation of force fields and water models in addressing IDP-specific studies [15, 22, 26, 30–35]. The core issue is that atomistic modeling of IDPs is inherently more demanding compared to globular proteins in their native states: IDPs exhibit multiple interconverting conformations with substantial populations.

In recent times, extensive studies have been done till date to address the prevailing challenges related to reproducing IDP conformational propensity with existing conventional force fields and water model combinations [36–38]. The case-specific systems varied from short dipeptides, IDP fragments, whole sequences of folded and unfolded IDPs. Nevertheless, the challenges in choosing appropriate (IDP-specific) potential energy functions to quantitatively reproduce corresponding structural/conformational propensity and corresponding molecular interactions, interactions persist [36]. Moreover, the solvation dynamics, associated with and important for biomolecular interactions, have not been adequately investigated till date, with changing force fields and water models. Further, the parameters defined for benchmarking the force-field performances were not uniform, and therefore lacking in identically reproducing experimental observable parameters. To overcome the aforementioned challenges, we evaluated the performance of thirteen combinations of biomolecular force field-water models (FF-wm, Table 1), in accurately representing the conformational ensembles of IDPs in extended unbiased MD simulations. We have investigated four representative IDPs: amyloid-*β*42 (A*β*42), a 42-residue peptide (PDB:1IYT) [39], Tau43 (PHF43) a 43-residue domain [40], amylin (human islet amyloid polypeptide (hIAPP)) a 37-residue peptide[41], and *α*-synuclein (*α*S) a 140-residue protein [42]. The chosen IDPs were selected based on their diversity: A*β*42, significant due to its aggregation properties and role in the pathology of Alzheimer’s disease (AD); Tau43, a 43 residue sequence derived from the 441-residue human Tau protein [43], identified as a key factor in the aggregation process that leads to AD [40]; 37 residue amylin (PDB: 2L86) [41], which is believed to be responsible for pathological progress of Type-II diabetes, and *α*S (PDB: 1XQ8) [42]an IDP with a larger sequence size and responsible for Parkinson’s disease (Fig.1). To verify globular conformational propensity, we simultaneously simulated two globular folded proteins of similar sequence size (HP36, the villin headpiece subdomain (PDB: 1VII)[44], and WW-domain (PDB: 1E0M) [45]). Our simulation protocols, inspired by previous studies [46], are unique in terms of reproducing diverse unstructured IDP phase-space (see Methods, section 4, Figure S1). The simulated data were then compared with experimental J-coupling parameters, which showed good agreement with the selected FF-wm combinations. Further, conformational propensity and water dynamics analyses provided a universal picture, as to which FF-wm serves best for IDPs in accurately reproducing structural and dynamic parameters.

**Fig. 1:**
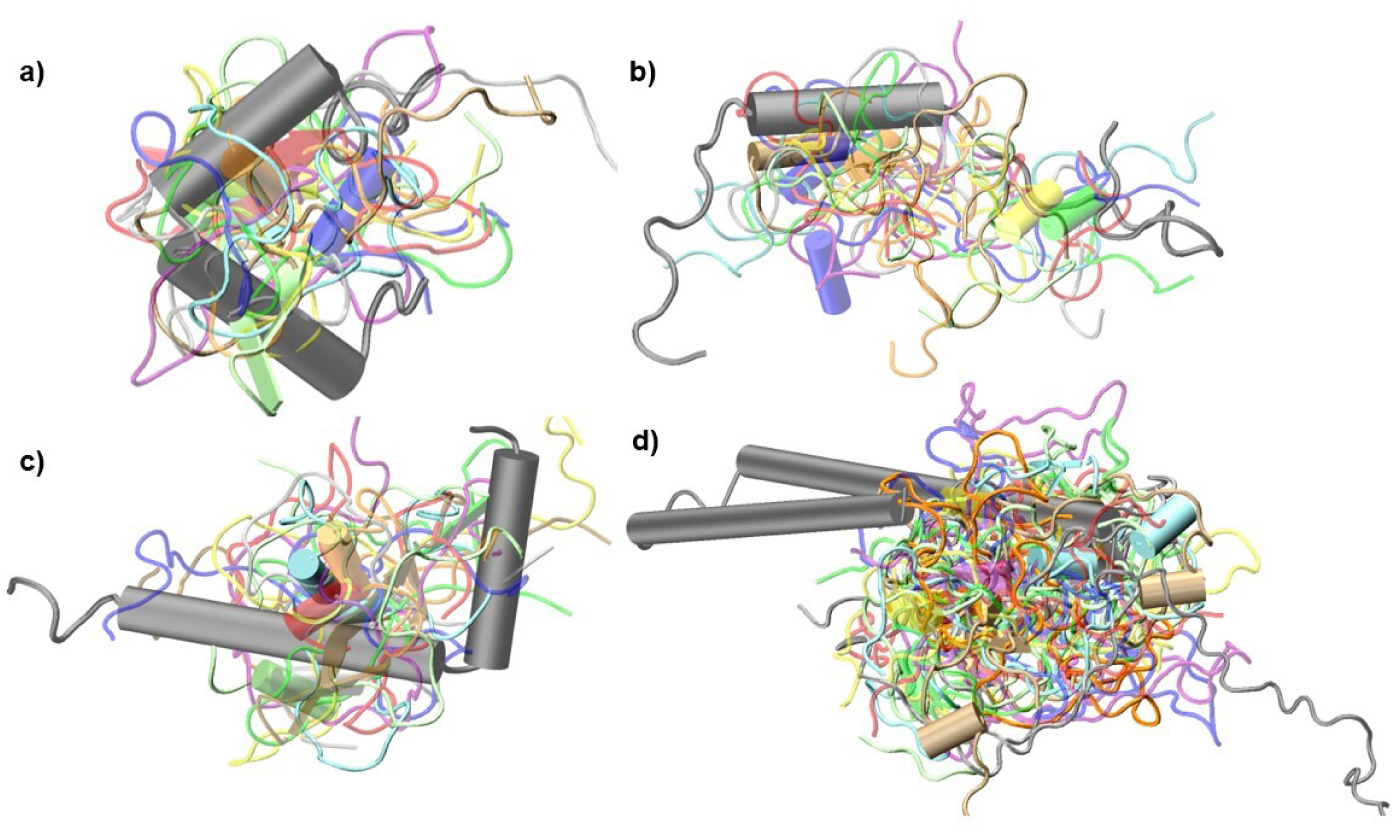
Representative snapshots of a) A*β*42, b) Tau43, c) Amylin, and d) *α*S. The helices are represented with cylindrical barrels, *β*-sheets with arrows. These are representative conformations from 10 independent simulations (see methods), superimposed on the initial PDB structure (denoted in gray), after removing the translational motion followed by a non-linear least square fitting.

**Table 1:** Force fields and water models.

| Serial Number | Force Field | Water Model |
| --- | --- | --- |
| FF-wm1 | A99SB-ILDN [47] | TIP4P-Ew [48] |
| FF-wm2 | A14SB [49] | TIP3P [23] |
| FF-wm3 | A99SB-ILDN-Dyes [50] | TIP4P-Ew [48] |
| FF-wm4 | C36-IDPSFF [51] | A99SB-disp [36] |
| FF-wm5 | C36-IDPSFF [51] | TIP4P-Ew [48] |
| FF-wm6 | CHARMM36m [33] | CHARMM TIP3P [52] |
| FF-wm7 | CHARMM36m [33] | TIP3P [23] |
| FF-wm8 | A14SB [49] | TIP4P-Ew [48] |
| FF-wm9 | CHARMM27 [53, 54] | CHARMM TIP3P [52] |
| FF-wm10 | A99SB [55] | OPC [56] |
| FF-wm11 | A99SB-disp [36] | A99SB-disp [36] |
| FF-wm12 | CHARMM27 [53, 54] | TIP3P [23] |
| FF-wm13 | GROMOS 96 (54a7) [57] | SPC/E [58] |

## 2 Results

By simulating the chosen IDPs in each of the FF-wm combinations (Table 1) we examined the IDP-specific effect of the chosen force-field water model combinations. Extended conformations are commonly employed as starting structures for IDP simulations as they facilitate conformational transitions in these disordered proteins. However, in the present work, our simulation protocol was designed to facilitate the transition from folded to unfolded states. (section 4, Figure S1). The generated conformational space for each IDP (specific to each FF-wm) was validated against previously obtained experimental observables (Tables 2, S2, S3, Fig. S2, S3). All calculated ^3^J*_HN_*_−_*_Hα_*-Coupling data are tabulated in SI (section S4). 2D density plots between the structural parameters (RMSD, Rg, SASA, end-to-end distance (E2E), structural persistence (P) [59]; see section S1) were analyzed and presented (Fig.2, S5, S6). In most of the cases, we observed noticeably broader distribution of values at 310 K compared with 300 K.

**Fig. 2:**
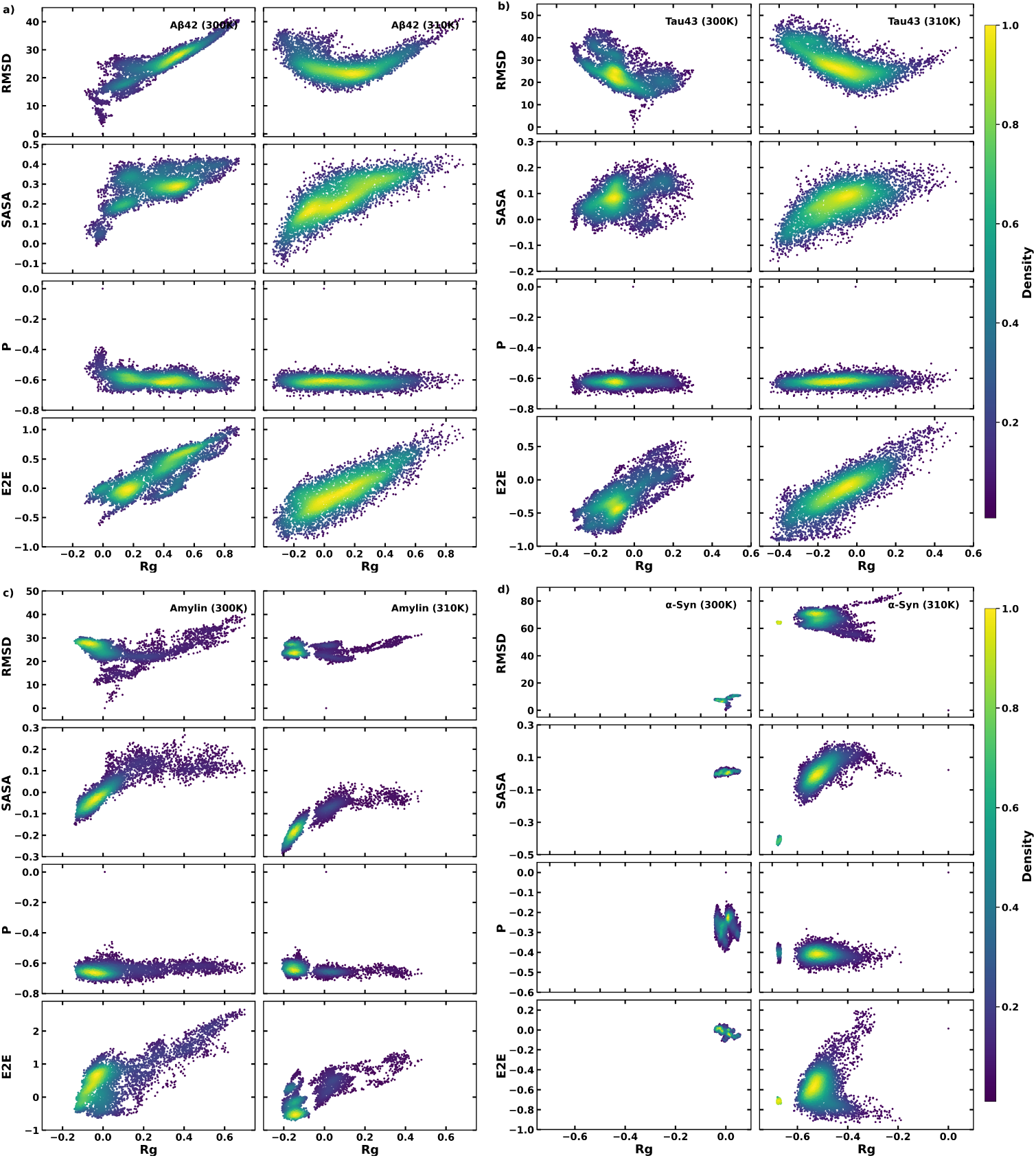
The 2D Kernel density plots as a function of Rg vs RMSD, Rg vs SASA, Rg vs P and Rg vs E2E as the coordinates for four selected IDPs: a) A*β*42, b) tau43, c) amylin and d) *α*S for FF-wm4.

### 2.1 Comparison with experimental observables

The NMR ^3^J*_HN_*_−_*_Hα_*-Coupling parameter is one of the robust experimental observables to test the performance of force fields for IDPs. The ^3^J*_HN_*_−_*_Hα_*-Coupling constants for the backbone *ϕ* angles were calculated using the gmx_chi gromacs module. It utilizes the Karplus equation [60] described as,

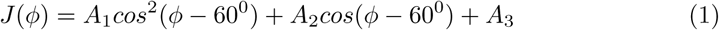

Where, J is the ^3^J*_HN_*_−_*_Hα_* coupling constant, *ϕ* is the backbone dihedral angle, and A1, A2, A3 are the empirically derived parameters. The gmx_chi code utilizes the parameters established previously [61] in which A_1_ = 6.51, A_2_ = - 1.76, and A_3_ = 1.60. ^3^J*_HN_*_−_*_Hα_* coupling data corresponding to each IDP for every FF-wm combinations, were obtained by averaging over ten independent trajectories (section 4, Fig. S1), sufficient for reproducing converged results for IDPs [51].

To quantitatively compare the ^3^J*_HN_*_−_*_Hα_* coupling data obtained from our simulations, with those obtained from experiments [43, 62, 63] and the predicted observables (*RC_3JHNHa Server*) [64], we employed four statistical metrics. These metrics are the cosine of the angle (cos *θ*) between the straight line (*y* = *x*) and the regression line corresponding to the correlation between experimental and calculated values, the Mean Fractional Deviation (MFD; Eq.2), the Mean Squared Error (MSE; Eq.3), and the Mean Absolute Error (MAE; Eq. 4).

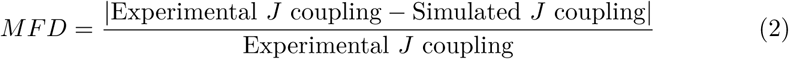

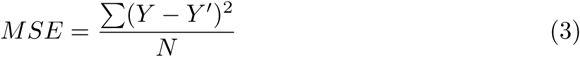

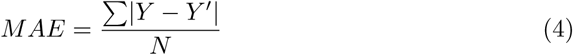

In the above equations, Y represents the simulated value of the ^3^*J_HN_*_−_*_Hα_*-Coupling, Y^′^ denotes the corresponding values from (*experiments* and/or *RC_3JHNHa Server*), N is the total number of ^3^J*_HN_*_−_*_Hα_*-Coupling data points included in the plot. The results for the statistical metrics are provided in MFD (Table 2), *cosθ* (Table S1), MSE (Table S2), and MAE (Table S3). A *cosθ* value of 1 reflects perfect agreement between the calculated and reference ^3^J*_HN_*_−_*_Hα_* data. Similarly, smaller deviation or error values indicate closer alignment between the simulation and experimental data. For A*β*42, the *cosθ* values range from 0.87 to 0.96 when fitted with experimental data and from 0.80 to 0.92 when fitted with the *RC_3JHNHa Server* data, with the highest agreement observed for FF-wm3. Tau43 shows more variability, ranging from 0.51 to 0.73 (when compared with experimental values) and 0.48 to 0.88 (in comparison with *RC_3JHNHa Server* predicted values). It is to be noted here that Tau43, by itself, is not a continuous sequence, but a combination of different segments of Tau441, coming together in brains affected by neurodegenerative diseases.(see 1). Since, NMR observables will have an environmental effect, the current calculated parameters, when compared with those obtained from the whole sequence Tau441 experimental observables, will therefore deviate more. The amylin system exhibits high agreement with *RC_3JHNHa Server* values, showing the highest alignment with calculated J-coupling values for FF-wm4, which lies between 0.83 and 0.94. Lastly, *α*S shows *cosθ* ranging from 0.55 to 0.96 (for experiment) and 0.54 to 0.94 (for *RC_3JHNHa Server*), with FF-wm4 and FF-wm5 showing the closest proximity with experimentally reported coupling values.

**Table 2:**
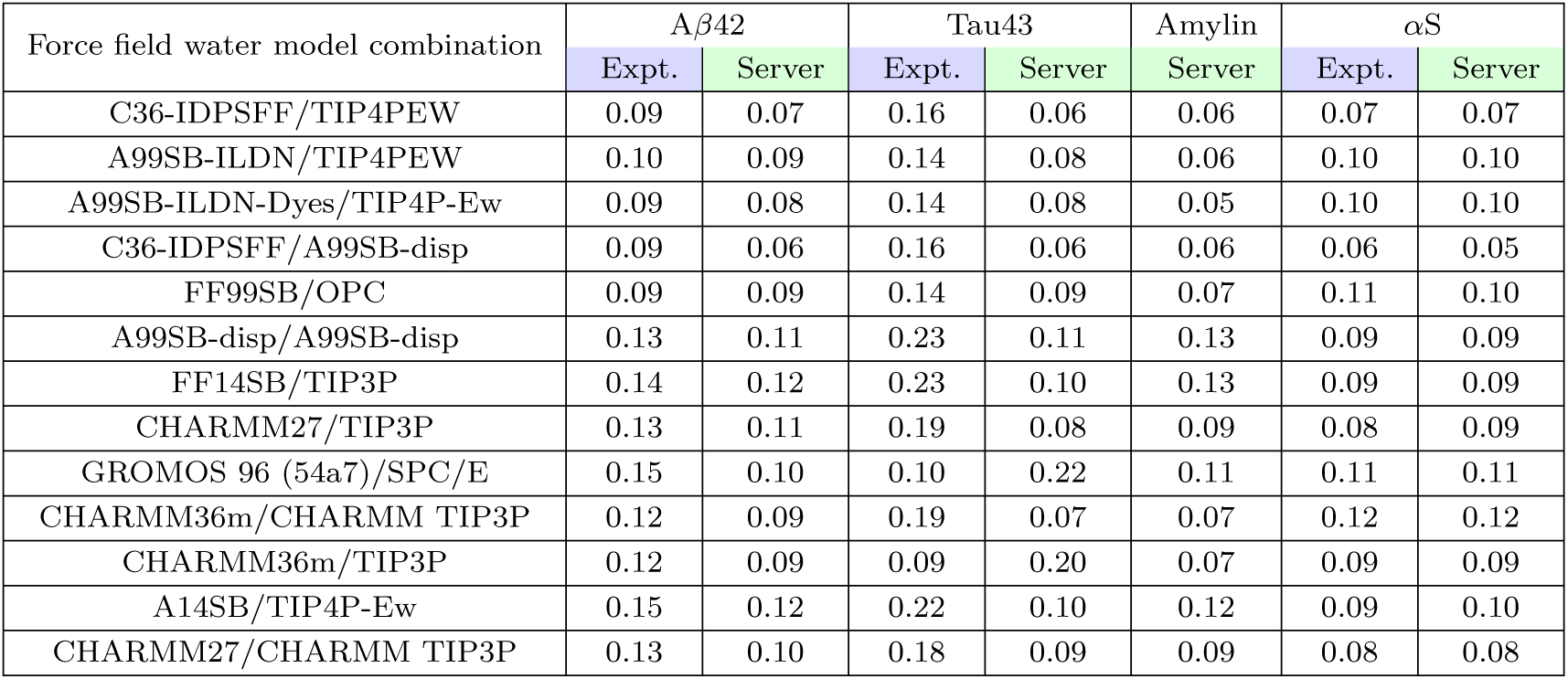
Mean fractional deviation (MFD) between experimental/*RC_3JHNHa Server* and calculated ^3^J*_HN_*_−*Hα*_-coupling constants of IDPs.

| Force field water model combination | $A\beta 42$ | | Tau43 | | Amylin | αS | |
| --- | --- | --- | --- | --- | --- | --- | --- |
|  | Expt. | Server | Expt. | Server | Server | Expt. | Server |
| C36-IDPSFF/TIP4PEW | 0.09 | 0.07 | 0.16 | 0.06 | 0.06 | 0.07 | 0.07 |
| A99SB-ILDN/TIP4PEW | 0.10 | 0.09 | 0.14 | 0.08 | 0.06 | 0.10 | 0.10 |
| A99SB-ILDN-Dyes/TIP4P-Ew | 0.09 | 0.08 | 0.14 | 0.08 | 0.05 | 0.10 | 0.10 |
| C36-IDPSFF/A99SB-disp | 0.09 | 0.06 | 0.16 | 0.06 | 0.06 | 0.06 | 0.05 |
| FF99SB/OPC | 0.09 | 0.09 | 0.14 | 0.09 | 0.07 | 0.11 | 0.10 |
| A99SB-disp/A99SB-disp | 0.13 | 0.11 | 0.23 | 0.11 | 0.13 | 0.09 | 0.09 |
| FF14SB/TIP3P | 0.14 | 0.12 | 0.23 | 0.10 | 0.13 | 0.09 | 0.09 |
| CHARMM27/TIP3P | 0.13 | 0.11 | 0.19 | 0.08 | 0.09 | 0.08 | 0.09 |
| GROMOS 96 (54a7)/SPC/E | 0.15 | 0.10 | 0.10 | 0.22 | 0.11 | 0.11 | 0.11 |
| CHARMM36m/CHARMM TIP3P | 0.12 | 0.09 | 0.19 | 0.07 | 0.07 | 0.12 | 0.12 |
| CHARMM36m/TIP3P | 0.12 | 0.09 | 0.09 | 0.20 | 0.07 | 0.09 | 0.09 |
| A14SB/TIP4P-Ew | 0.15 | 0.12 | 0.22 | 0.10 | 0.12 | 0.09 | 0.10 |
| CHARMM27/CHARMM TIP3P | 0.13 | 0.10 | 0.18 | 0.09 | 0.09 | 0.08 | 0.08 |

Mean fractional deviation (MFD), represents the average of residual fractional deviation between the simulated data and those obtained from the experiment and/or *RC_3JHNHa Server*. For A*β*42, the MFD ranges from 0.15-0.09, and the lowest fractional deviation is observed for FF-wm5, FF-wm3, FF-wm4, and FF-wm10. When fitted with the *RC_3JHNHa Server*, the values lie in the range 0.06-0.12, with FF-wm5 and FF-wm4 showing the highest agreement, with both *RC_3JHNHa Server* as well as with the experimental data. In the case of Tau43, values lie between 0.09 and 0.23 when fitted with experiments, and 0.06-0.23 when fitted with the server. The values are more variable, with maximum concordance with the server-generated J-coupling values, observed for the FF-wm5, FF-wm3, and FF-wm4. For amylin, values lie between 0.05 and 0.19, with the highest agreement for FF-wm3 followed by FF-wm5, FF-wm1, and FF-wm4. Similarly, *α*S shows the optimal agreement between the experimental / *RC_3JHNHa Server*-data and simulated data for FF-wm4. A similar pattern could be observed for the MSE and MAE with the closest alignment with the reference coupling values, and those from FF-wm4.

Based on the value of cos *θ* (Table S1), the force field–water model combinations, FF-wm5, FF-wm1, FF-wm3, FF-wm4, and FF-wm10 force field-water model combinations demonstrate the best performance. Looking at the error values, MFD (Table 2), MSE (Table S2) and MAE (Table S3), the lowest error values are observed for the FF-wm4 followed up with the FF-wm5 while others have marginally pronounced error values and are close to each other.

### 2.2 Structural Propensity of IDPs

The structural dynamics of IDPs were assessed under selected sets of four FF-wm combinations (Table 1), based on NMR J-coupling validations. We have analyzed the fractional values of the key metrics root mean square deviation (RMSD), radius of gyration (Rg), solvent accessible surface area (SASA), end-to-end distance (E2E), and P (secondary structure persistence parameter [59]) for the chosen FF-wm combinations. The fractional values are dimensionless quantities, computed w.r.t. reference structure (for equations see SI section S2). The negative values indicates a lesser value of that parameter w.r.t. the reference structure. These parameters provide insights into conformational variability, compactness, and structural integrity under thermal fluctuations at 300 K and 310 K [65]. RMSD quantifies the change in conformation with respect to a reference structure (see Section S1.1), P shows the change in secondary structural propensity with respect to a reference structure (see SI, Section S1.3). The density plot in Fig. 2 depicts the relationship between these structural features, providing a comparative view of IDP behavior with the varying combinations of FF-wm. It is evident that the RMSD distribution exhibits an overall increase at 310 K for most IDPs, signifying greater conformational diversity at elevated temperatures. Specifically, for A*β*42, the RMSD density is concentrated around 30 at 300 K but shifts to 25 at 310 K while maintaining a broader spread toward lower values at 300 K, suggesting enhanced structural heterogeneity at the lower temperature. For SASA, a general shift toward lower values is observed for most FF-wm combinations (Fig. S4, S5, S6), except for FF-wm4, where two out of the four IDPs display a relatively higher SASA distribution at 310 K. This suggests that FF-wm4 promotes a more solvent-exposed ensemble compared to the other three FF-wm combinations, which exhibit greater compaction. Interestingly, the E2E distribution does not exhibit a consistent trend as a function of temperature. However, IDPs tend to show lower E2E distances at 310 K than at 300 K, suggesting that their terminal regions might come closer together. This behavior aligns with the observed reduction in Rg at 310 K, further supporting the hypothesis of a molten-globule-like conformational propensity at higher temperatures[66].

**Table 3:**
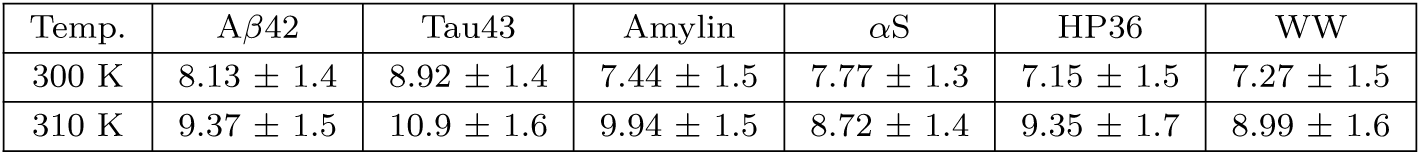
Diffusion coefficient (×10^−4^Å^2^*/*ps) of water molecules around proteins (Sh_1_). The table reports the average diffusion coefficients (from every 2 ps time-window, from 2-20 ps MSD data) and their standard deviations in round brackets for FF-wm4.

| Temp. | Aβ42 | Tau43 | Amylin | αS | HP36 | WW |
| --- | --- | --- | --- | --- | --- | --- |
| 300 K | $8.13 \pm 1.4$ | $8.92 \pm 1.4$ | $7.44 \pm 1.5$ | $7.77 \pm 1.3$ | $7.15 \pm 1.5$ | $7.27 \pm 1.5$ |
| 310 K | $9.37 \pm 1.5$ | $10.9 \pm 1.6$ | $9.94 \pm 1.5$ | $8.72 \pm 1.4$ | $9.35 \pm 1.7$ | $8.99 \pm 1.6$ |

Furthermore, the secondary structure persistence parameter (P) consistently shows lower values at 310 K across all IDPs, indicating a loss of secondary structural content. This reduction, combined with lower E2E and Rg values, suggests a transition toward a disordered structures, resembling the corresponding characteristics often observed in partially folded or molten-globule states. Notably, FF-wm4 promotes relatively extended conformations compared to other FF-wm combinations, and also replicating near experimental observables that indicate a preference for expanded states in specific IDPs.

### 2.3 Water Dynamics

Water molecules play a significant role in reshaping and influencing the structural and functional characteristics of proteins and IDPs [67–69]. IDPs rely heavily on their hydration environment to maintain their functional disorder [69]. The study of water dynamics, which examines the organization and movement of water molecules around these IDPs, provides critical insights into the molecular-level details of biological processes [67, 70–72].

Previous studies on validating force-fields and water models have mostly underscored the structural propensity and corresponding resemblance with experimental results.

Our current studies have identified four best FF-wm combinations in terms of experimental validation and structural propensity (see Table 2, section 2.1 and Section 2.2) for IDPs.

#### 2.3.1 Radial Distribution Function

Initially, we have identified the solvation shells by calculating the radial distribution function g(r) between water oxygen (OW) and protein-heavy atoms (Fig 3). The g(r) is defined as:

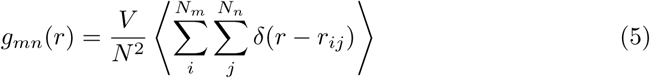

here, r*_ij_* is the distance between protein-heavy atoms and water-OW, *V* is the volume and *N* is the total number of atoms, *N_m_* and *N_n_* are the total number of water oxygen and protein-heavy atoms respectively.

**Fig. 3:**
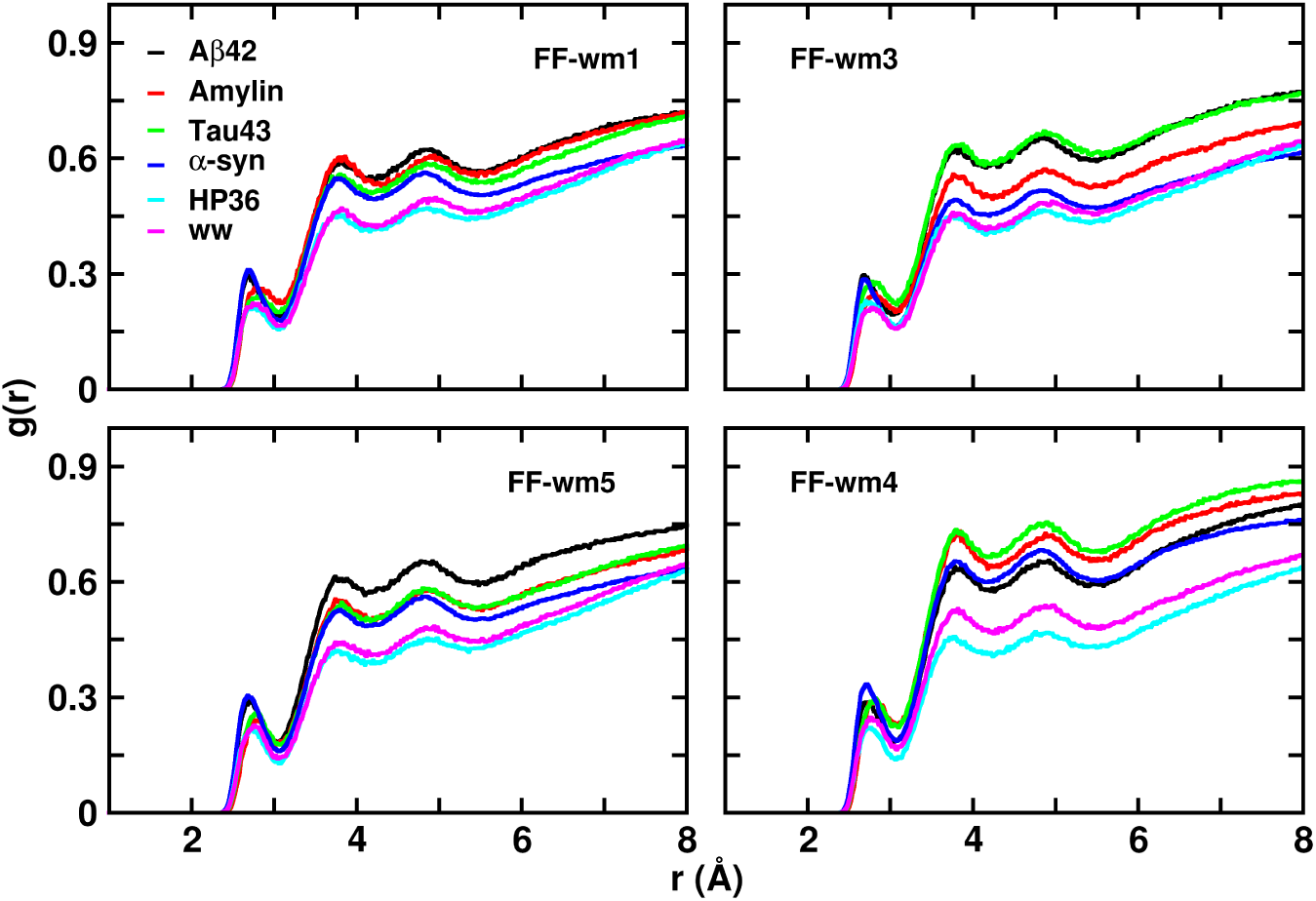
Radial distribution function (g*_mn_*(r)) of OW-water with protein heavy atoms, for the chosen FF-wm combinations at 310 K. Data corresponding to HP36 and WW domain are from 300 K simulations.

The height and position of the peak defines the hydration shells and offer insights into the structural ordering of water molecules in the solvation layer. The first peaks at 3.06 Å, represents the first hydration shell (Sh_1_) where water molecules form strong hydrogen bonds or interact electrostatically with the protein [73]. The second hydration shell (Sh_2_) is at 4.2 Å, where water molecules are less structured but still influenced by the presence of the protein. We also defined the third hydration shell (Sh_3_) at 5.5 Å, and beyond this (∼ 5–8 Å), g(r) gradually approaches near bulk-like water behavior (Sh_4_) [73]. All the four IDPs show prominent peaks at Sh_1_, depicting a well-defined and ordered first hydration layer. The peaks corresponding to Sh_2_, and Sh_3_, are also prominent.

#### 2.3.2 Mean Squared Displacement

Based on the defined hydration shells (section 2.3.1), we have calculated the mean squared displacement (MSD) of water molecules, which provides insights into the displacement of water molecules and alterations in their mobility due to water-protein interactions. Single-particle dynamics of water molecules can be characterized through MSD:

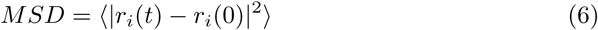

here, *r_i_*(0) and *r_i_*(t) are the coordinates of the *i^th^* molecules at time *t* = 0 and *t* = *t* respectively, and ⟨ *….*⟩ is the ensemble average. We further evaluated the self-diffusion coefficient, *D*, using the Einstein relation by linear regression fitting over the MSD data (from 2-20 ps averaged over every 2 ps time-window) using the equation:

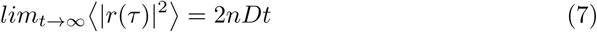

here *n* is the dimensionality of the system, *D* is diffusion coefficient and *t* is time.

Unlike globular proteins, IDPs have a higher degree of surface exposure to water. Therefore, dynamic studies of the hydration water helps to elucidate the dependence of hydration water on the solute (protein) conformation, providing a unique perspective on the behavior of IDPs in solution. Based on the structural parameters, we have evaluated the FF-wm combinations in reproducing the extensivity and heterogeneous conformational propensity of IDPs, in accordance with experimentally determined NMR observables. To the best of our knowledge, this is the first quantification and comparison of the effect of chosen FF-wm(s) in the surrounding aqueous environment. We already defined the hydration shells (Sh_1_, Sh_2_, Sh_3_, Sh_4_, see section 2.3.1, Fig.3)). The calculated (shell-specific) MSD are plotted in Fig. 4, S7 (corresponding D values are tabulated in Tables S14, S15, S16, in SI section S6). From Fig. 4, the MSD is lowest in solvation shell Sh_1_, indicating highly restricted dynamics due to strong interactions with IDP-surface. The MSD increases gradually with distance from the protein surface, it shows the reduced influence of the protein residue on the corresponding translational dynamics of water molecules in respective shells. The bulk water shows highest MSD, depicting unrestricted water dynamics for all the water models studied. At 310 K, MSDs for all the shells are modestly higher than those at 300 K, indicating enhanced water mobility due to increased thermal energy. This increase in mobility is more notable for the outer hydration layers (Sh_3_, Sh_4_) and bulk water compared to Sh_1_. In the latter, interactions with the IDPs remain dominant. We further observed similar trends for A*β*42 and *α*S across the solvation shells. The differences in MSD between shells are less pronounced for *α*S compared to A*β*42, suggesting weaker coupling between water and *α*S. A*β*42 shows a more pronounced restriction in the solvation dynamics of Sh_1_ and Sh_2_, compared to *α*S, likely due to stronger or more structured interactions between water and A*β*42. The conformational propensity of *α*S is more heterogeneous and relatively more collapsed, reflected from a decrease in Rg and increase in RMSD distribution, compared to its initial dimension, as reported above (Fig. 1, Fig. 2). Previous studies have also underscored the conformational heterogeneity of *α*S [74]. The molten globule like, yet disordered conformations for *α*S, obtained in our current study, is plausibly leading to the less protein-water interactions. We have also observed consistent dynamic stratification in the hydration shells of Tau43 and Amylin, with water mobility progressively increasing from the protein surface outward, similar to A*β*42 and *α*S. In both Tau43 and Amylin (Fig. S7), bulk water shows the highest MSD, while the innermost shell (Sh_1_) shows the lowest, indicating strong protein-water interactions near the surface. The effect of temperature is also preserved here, with higher MSDs at 310 K than 300 K, especially for outer shells and bulk. We have also observed that across the chosen FF-wm combinations, FF-wm4 shows lower MSD values in Sh_1_ and Sh_2_, which suggests stronger water-protein interactions, likely due to the dispersion-enhanced water model. The A99sb-disp water allows for more structured hydration shells around solutes due to its fine-tuned interaction potential. Other FF-wm combinations (Fig. 4 b,c,d) have higher MSD values, which show less structured hydration shells compared to A99sb-disp water. The TIP4P-Ew water model may not model dispersion effects as strongly, leading to weaker coupling with the protein surface.

**Fig. 4:**
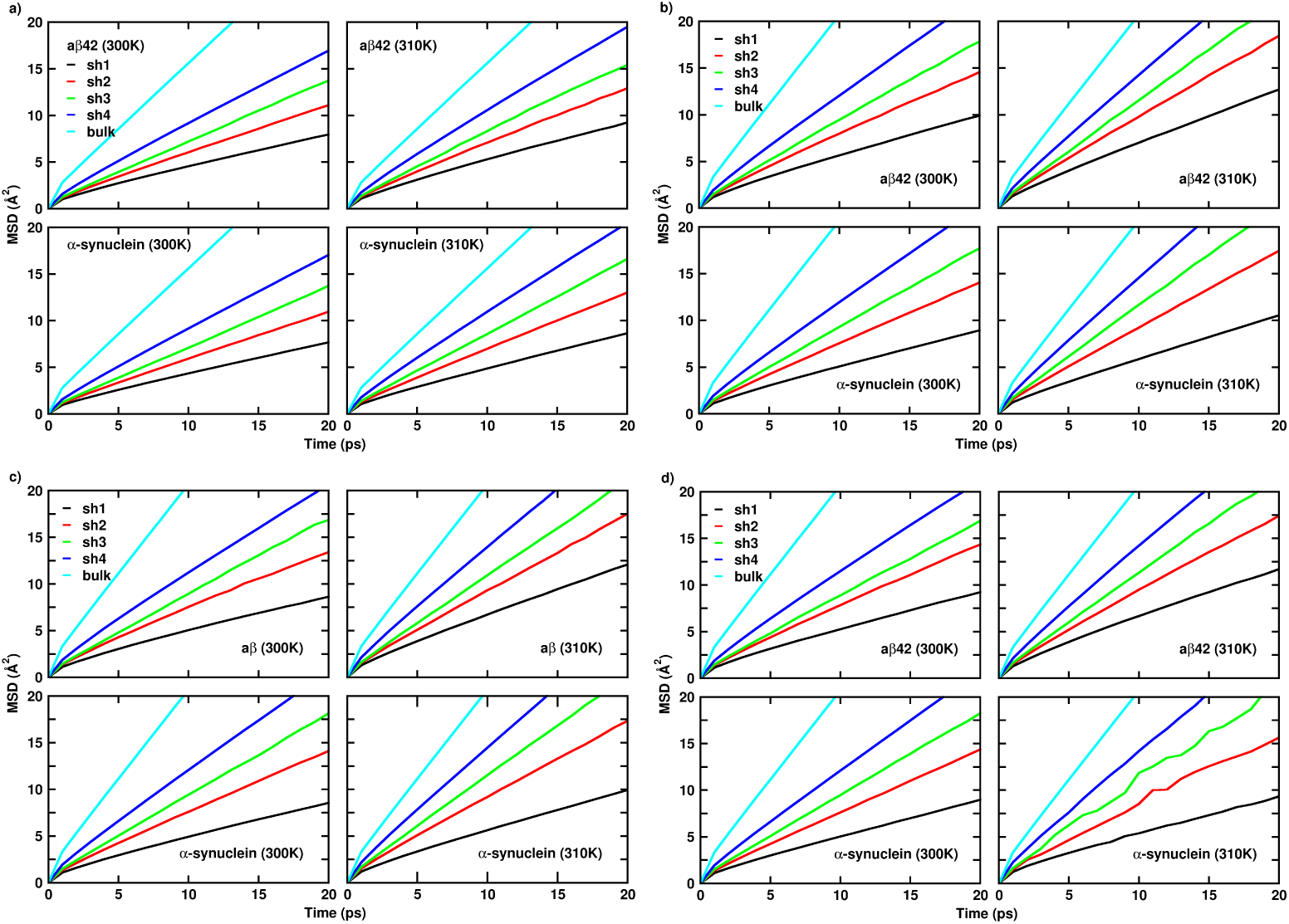
Mean squared displacement (MSD) of the hydration shells (Sh_1_, Sh_2_, Sh_3_, and Sh_4_) surrounding the proteins (A*β*42 and *α*S)at temperatures 300 K and 310 K, based on the hydration layer peaks identified using g(r) (see Fig.3) for a)FF-wm4, b)FF-wm5, c)FF-wm3 and d)FF-wm1. Bulk water (without protein) data is added for comparison.

The outer hydration shells (Sh_3_, Sh_4_) in all the combinations for all the systems (Fig. 4, S7), show a gradual increase in MSD values, reflecting the transition to bulk water-like dynamics. However, A99sb-disp water (Fig.4, a) shows a more pronounced deviation from bulk behavior even in outer shells, likely due to its ability to sustain longer-range interactions. TIP4P-Ew (Fig.4 b, c, d) converge faster to bulk dynamics, indicating less persistent hydration effects. At 300 K, the remaining FF-wm combinations show restricted water dynamics in inner shells, but the restriction is most pronounced in FF-wm4, followed by FF-wm5. The calculated MSD in shells Sh_1_ and Sh_2_ with FF-wm3 and FF-wm1, suggest less tight coupling with water. At 310 K, all combinations show an increase in MSD values, with the effect being more significant for TIP4P-Ew water models. This reflects a temperature-dependent loosening of water-protein interactions, and faster water dynamics (Fig.4 b, c, d). Bulk water dynamics of A99sb-disp water model (Fig.4 a) shows slightly lower MSD trend indicating a more structured water model compared to TIP4P-Ew (Fig.4 b, c, d). Although, the dispersion interactions are apparently heightened for hydration shells surrounding globular proteins HP36 and WW-domain (Fig. S8), our simulations exhibit protein-water dispersive interactions at distal sights from the surface, even for the unstructured IDPs, with FF-wm4.

Earlier studies reported highly restricted and anomalous diffusion of water molecules near protein surfaces [75]. However, regions near IDPs exhibited comparatively lesser retardation in translational motion, attributed to a higher number of water–water hydrogen bonds and tetrahedral arrangements than those observed near ordered/globular proteins. [76, 77]. In similar lines, to quantify the differences in MSD for the respective systems, we have defined an additional parameter D1_diff_ (See SI section S6.1). From Fig. 5, we see that for FF-wm4, D1_diff_ is less separated from each other, for A*β*42 and Amylin. Quantification of D1_diff_ for Tau43 and *α*S show relatively more differences between the chosen shells. On the other hand, for another representative FF-wm combination (FF-wm5, Fig. S9), D1_diff_ is in general, widely separated between the chosen hydration shells. We already found that FF-wm4 combination has an increased effect on the solvation layer surrounding IDPs, due to the presence of a99sb-disp water model (Fig. 4). This is not observed for another case-specific FF-wm combination (Fig. S9). D1_diff_ of our chosen globular proteins, exhibits increased dispersion effect in the hydration layer at distal sites from the protein surface, only for the WW-domain, with FF-wm4 combination (see Fig. S10).

**Fig. 5:**
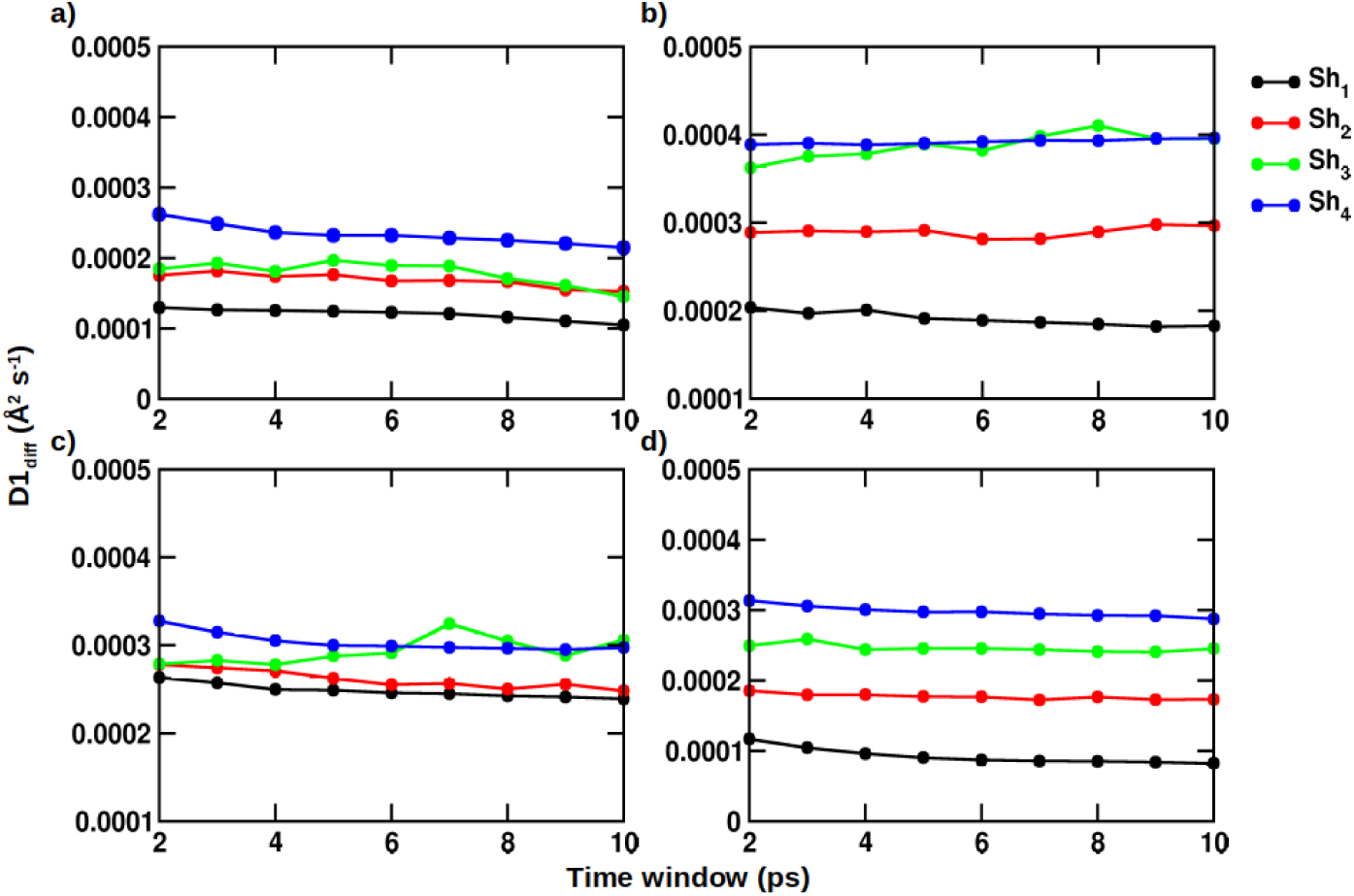
Differences in the mean squared displacement for the respective hydration shells, at the different time domains (see Table 3), as a difference of the two simulated temperatures, for a) A*β*42, b) Tau43, c) Amylin, d) *α*S, for FF-wm4 (Table 1).

We have also defined D2_diff_ (SI Section S6.1), to investigate the differential behavior of the respective dynamics of the hydration shells, for two case-specific FF-wm combinations (FF-wm4, FF-wm3). D2_diff_ (Fig. 6, S11) is found to be in accordance with temperature effects as mentioned before (see Section 2.3.2, second paragraph). Interestingly, when D2_diff_ is computed with Sh_3_, it is found to increase with time, and the difference between the two temperatures is more pronounced in comparison to Sh_1_. D2_diff_ further remains similar in general, when computed between the two adjacent shells (Sh_3_, Sh_2_; Sh_2_, Sh_1_). However, upon calculating D2_diff_ w.r.t. Sh_4_, we find a corresponding decay, when compared with the remaining hydration shells (Sh_3_, Sh_2_, Sh_1_). Sh_1_, being present in the closest proximity of the IDP surface, has the most restricted dynamic behavior. The restricted dynamical behavior decreases at 310 K when compared with 300 K, as evident from D2_diff_ trend for Sh_3_ - Sh_2_. This indicates water to be relatively more restricted at Sh_1_, at 310 K, compared to Sh_3_. However, the decay in trends of D2_diff_ involving Sh_4_ implies more intricate factors in regulating water dynamics at distal hydration layers from protein surfaces. These suggest that while the hydration shells around IDPs are highly sensitive to temperature and exhibit greater mobility at higher temperatures, the distance of hydration shells (proximal or distal) from IDPs also has a crucial role to play. While, for D1_diff_, the trend for IDPs and chosen folded proteins partly match considering FF-wm4; similar trends have been found for all the proteins, in case of D2_diff_ (Fig. S12), across the chosen FF-wm combinations. These allow for further intricate quantification of water dynamics at distal sites, surrounding biomolecular surfaces.

**Fig. 6:**
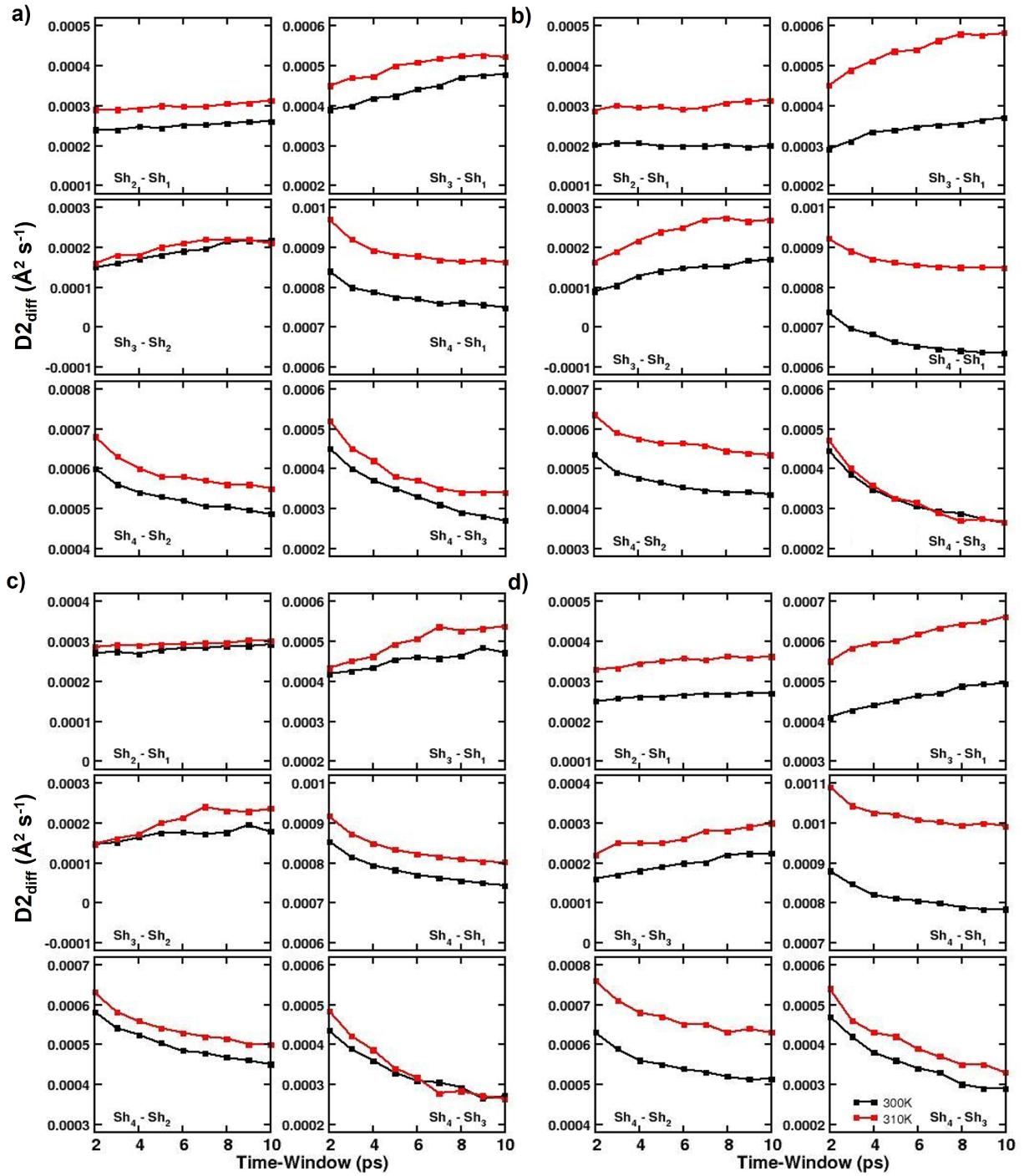
Differences in the mean squared displacement between the consecutive hydration shells (Fig.3), at the different time domains (see Table 3), for the two simulated temperatures, for a) A*β*42, b) Tau43, c) Amylin, d) *α*S, for FF-wm4 (Table 1).

## 3 Discussion and Conclusion

Several groups have performed significantly long simulations to test the accuracy of the existing force fields for IDPs [78]. Earlier studies have even shown the sensitivity of the force field-water model combination in calculating NMR observable [51, 79–81]. Reports identifying TIP4P-D being capable at improving the reliability of IDP simulations can be found in the literaure [79]. In the present study, an extended simulation of the folded and unfolded states for the chosen IDPs has been undertaken across thirteen FF-wm combinations. The NMR ^3^J*_HN_*_−_*_Hα_*-Coupling analysis demonstrates that the FF-wm4 force field-water model combination consistently provides the best agreement with experimental and *RC_3JHNHa Server*-predicted coupling constants across various IDPs, imminently reflected by high cos *θ* values and low error metrics (MFD, MSE, MAE). FF-wm5 also shows strong performance, while other combinations yield slightly higher deviations but remain comparable. A total of four FF-wm combinations show good agreement when compared with the experimental parameters. Based on the NMR ^3^J*_HN_*_−_*_Hα_*-Coupling analyses, we further tried to see whether these FF-wm combinations generate conformational phase-space, in accordance with experimental observations. The structural propensity of IDPs was analyzed across different force field-water model (FF-wm) combinations using metrics such as RMSD, Rg, SASA, E2E, and P. The results revealed that RMSD distributions were generally higher at 310 K for all chosen IDPs, indicating greater structural diversity. SASA distributions showed lower values for most FF-wm combinations at 310 K, except for FF-wm4, which exhibited relatively higher SASA for two IDPs. E2E distributions did not follow a specific temperature-dependent pattern but were generally lower at 310 K, suggesting a tendency for head-tail group interactions. A decrease in P-value at 310 K indicates a loss in secondary structure, leading to molten-globule-like conformations. Among the FF-wm combinations, FF-wm4 relatively produced more extended conformations that closely align with experimental observations, highlighting its superior performance in replicating IDP behaviors.

Previous studies [73] have already distinguished the different hydration shells depending on the dynamic nature of water molecules (also underscored in Fig.3). The noticeable information here lies in the fact that the hydration shells are well-defined across all IDPs as well as for the chosen globular proteins, irrespective of the different FF-wm combinations. We have therefore studied the crucial role of hydration water dynamics in understanding IDP behavior, with strong dependence on the force field-water model (FF-wm) combinations. Among the combinations, FF-wm4, featuring the A99sb-disp water model, exhibits the strongest coupling between water and IDPs, as indicated by lower MSD values in the inner hydration shells (Sh_1_, Sh_2_) and more structured hydration effects. This enhanced interaction is attributed to the dispersion effects of the A99sb-disp water model, which propagate to outer shells and deviate from bulk water dynamics. Comparatively, other water models like TIP4P-Ew show faster convergence to bulk-like behavior, indicating weaker hydration effects. Temperature significantly influences hydration dynamics, with higher mobility observed at 310 K, particularly in outer shells (Sh_3_, Sh_4_). The differential diffusion analysis (*D*_diff_) further reveals distinct trends in hydration dynamics, emphasizing the pronounced restriction in inner shells at elevated temperatures. Overall, FF-wm4 provides the most acceptable representation of hydration dynamics around IDPs, capturing both structural and dynamic properties with anticipated understanding.

## 4 Methods

Four IDPs chosen for the present work are A*β*42 (PDB:1IYT) [39], amylin (PDB: 2L86) [41], *α*S (PDB: 1xq8) [42] and Tau43. The initial structure for the 43-residue sequence Tau43, derived from the 441-residue human Tau protein, was taken from previously reported work [40]. MD simulations of the selected four IDPs were performed with the GROMACS version 2021.4 patched with PLUMED [82],[83]. In this study, a set of different force fields, including GROMOS, CHARMM, and AMBER, was used, developed for IDPs and water models with three and four sites, respectively (these force fields are listed in Table 1). Each system is prepared by solvating a cubic box containing a protein at the center. Counter-ions were added to neutralize the system, depending on the total charge of each IDPs. The Verlet algorithm with an integration timestep of 2 fs was used to integrate the equations of motion. Bonds involving H-atoms were constrained using the LINCS algorithm [84]. A cutoff of 1.0 nm was applied for short-range electrostatic and Lennard-Jones interactions. Long-range electrostatic interactions were calculated using the Particle Mesh Ewald (PME) method with fourth-order interpolation. [85]. Dispersion correction was used to adjust the energy and pressure terms to compensate for the truncation of van der Waals interactions. Temperature was controlled with the Bussi-Donadio-Parrinello thermostat [86], with a relaxation time of 0.1 ps. Separate coupling groups were used for the protein and water/ions to couple with the temperature bath. Parrinello-Rahman barostat [87] was used to maintain the pressure fixed at 1 bar, with a relaxation time of 2 ps and isothermal compressibility of 4.5 *times*10^−5^ bar^−1^. Each system was subjected to energy minimization using the steepest descent algorithm. Initial equilibration run for each system was done under canonical (NVT) ensemble for 100 ps, followed by 100 ps isothermal-isobaric (NPT) ensemble equilibration. Positional restraints were applied on the IDPs at equilibration steps, which were later removed during the production run of 100 ns at 300 K. Following these, simulated annealing was performed to achieve the disordered ensembles of IDPs. All the systems were annealed with 7 annealing points with a starting temperature of 300 K, 350 K (300 ps), 400 K (300 ps), 450 K (300 ps), 500 K (300 ps), 550 K (300 ps), 600 K (500 ps). The final configurations of all the systems were subjected to another NVT (canonical ensemble) equilibration for 100 ns at 600 K, where the configurations were saved after every 10 ns to create 10 distinct sets. Each set was gradually cooled stepwise from 600 K down to 310 K, similarly using a simulated annealing technique. Systems were cooled down using 7 annealing points with a starting temperature of 600 K, 550 K (300 ps), 500 K (300 ps), 450 K (300 ps), 400 K (300 ps), 350 K (300 ps), 310 K (500 ps). All ten sets were subjected to 1 ns NPT equilibration at 310 K, with positional restraints on the protein-heavy atoms. This was followed by a trajectory generation in the NPT ensemble for 100 ns of resolution 10 ps for all ten sets, resulting in a total of 1 *µ*s trajectory generation for each IDP and force field-water model combination. The resultant sets were used for the conformational analysis of the proteins. For the water dynamics analysis, a separate trajectory of 1 ns in the NVT ensemble was generated at a resolution of 1 ps. A separate simulation for bulk water was performed using a water box containing 2048 water molecules and a molecular density of 33 molecules nm^−3^.

## Acknowledgements.

The authors acknowledge the support of IHub-Data and DST-SERB (CRG/2021/008036) for financial support and HPC at the International Institute of Information Technology (IIIT), Hyderabad for computational resources.

## Author Contributions

U.D.P. and P.C. conceived and supervised the study. V.K., A.S., and P.C. contributed to the writing of the manuscript, V.K., P.C., A.S., and M.M. wrote the codes and analyzed the data, V.K. formulated the simulation protocol, and V.K. and M.M., and A.S. contributed to the simulation. All authors approved the final version.

## Code and Data availability

The data and code used in this work can be accessed through the GitHub repository https://github.com/devalab/BenchIDPs.

## Supplementary Information

**Fig. S1:**
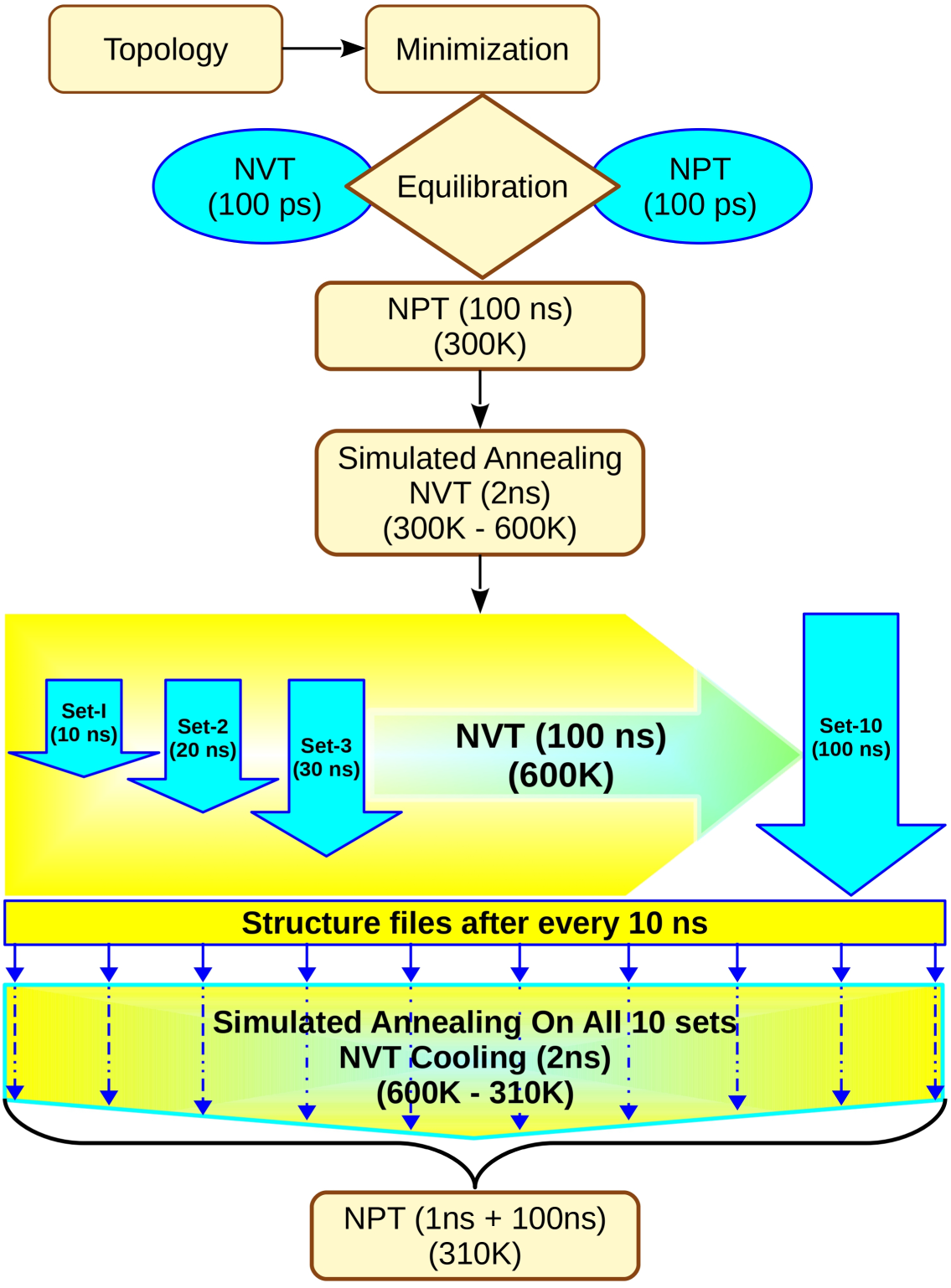
Flow Chart for the IDP simulation protocol.

### S1 Formula for measuring conformational properties

#### S1.1 RMSD

The Root Mean Square Deviation (RMSD) quantifies the deviation of atomic positions from a reference structure:

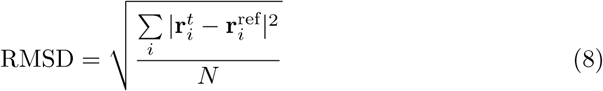

where N= number of atoms, 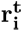-position of atom i in the current structure, 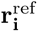= position of atom i in the reference structure.

#### S1.2 Rg

The radius of gyration (Rg) is a measure of the overall compactness of a protein structure:

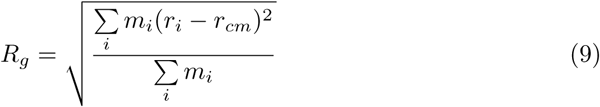

where,m*_i_*= mass of atom i, usually uniform for all atoms in simulations, r*_i_*= position vector of atom i, r*_cm_*= position vector of the center of mass.

#### S1.3 Structural Persistence (P)

The structural persistence P quantifies the extent of structural destabilization in proteins relative to a reference structure.

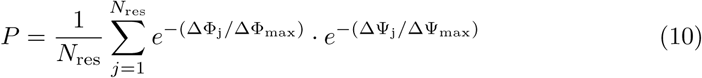

where,N_res_= Number of residues in the protein, ΔΦ*_j_* and ΔΨ*_j_*= absolute values of the changes in the dihedral angles of residue j over the reference values, ΔΦ*_max_* and ΔΨ*_max_*= Maximum alterations possible in the Ramachandran diagram.

#### S1.4 SASA

Solvent-accessible surface area (SASA) is a biomolecule’s surface area accessible to a solvent.

#### S1.5 E2E

The end-to-end (E2E) distance is the distance between the C*α* atoms of the N- and C-termini.

### S2 2D Density Plot Formula

#### S2.1 Fractional Root Mean Square Deviation

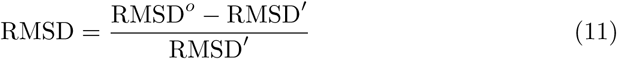

where,RMSD*^o^*= RMSD of each frame at production run (300 K or 310 K) w.r.t. the PDB structure RMSD*^′^* = RMSD of frame after equilibration w.r.t. the PDB structure.

#### S2.2 Fractional Radius of gyration

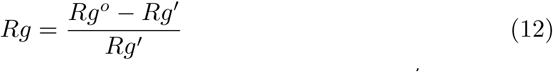

where, Rg*°*= Rg of each frame at production run (300 K or 310 K) Rg*^′^* = Rg of the PDB structure.

#### S2.3 Fractional Solvent Accessible Surface Area

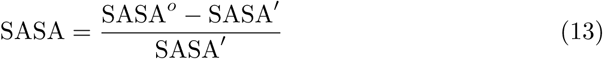

where, SASA*°*= SASA of each frame at production run (300 K or 310 K) SASA*^′^* = SASA of the PDB structure.

#### S2.4 Fractional End-to-End distance

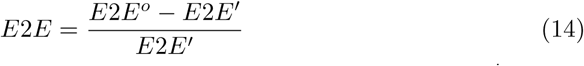

where, E2E*°*= E2E of each frame at production run (300 K or 310 K), E2E*^′^* = E2E of the PDB structure.

#### S2.5 Fractional Structural Persistence

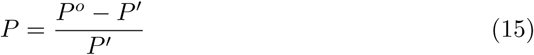

where, P*^o^*= P of each frame at production run (300 K or 310 K) w.r.t. the PDB structure, P*^′^* = P of frame after equilibration w.r.t. the PDB structure.

### S3 Comparison with experimental observable

**Fig. S2:**
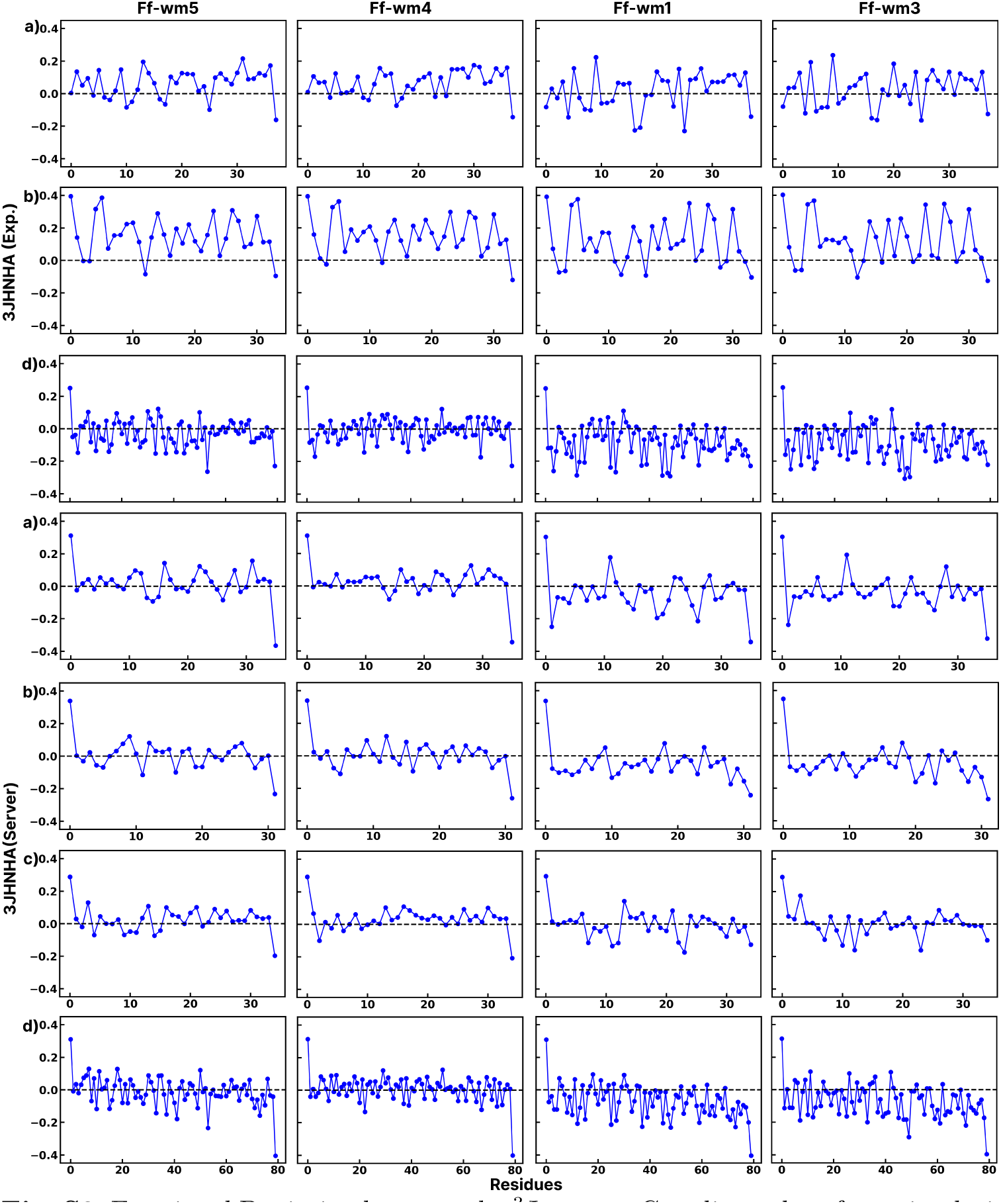
Fractional Deviation between the ^3^J*_HN_*_−_*_Hα_*-Coupling values from simulation and the NMR data from experiment/*RC_3JHNHa Server*. a) A*β*42, b) Tau43, c) amylin and d) *α*S

**Fig. S3:**
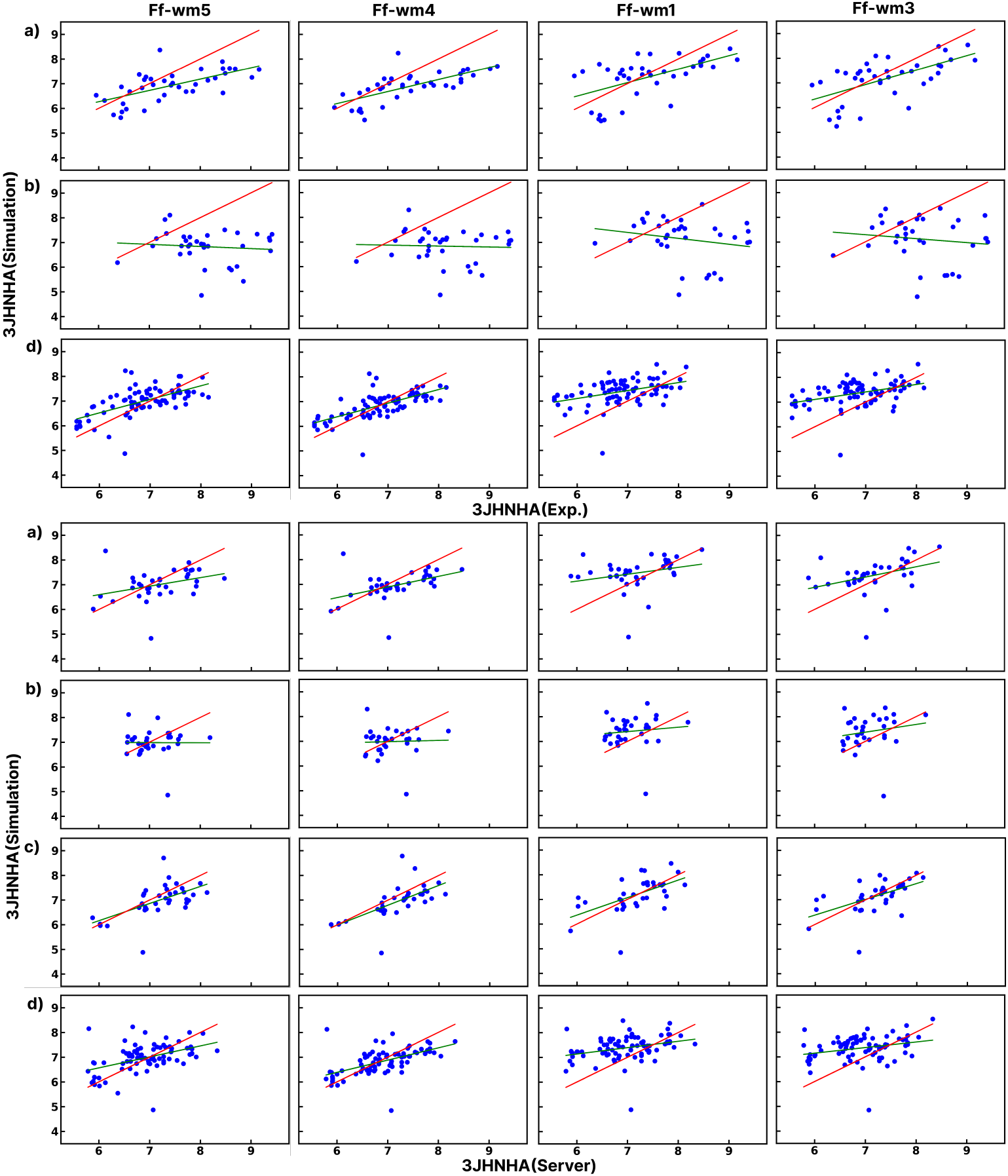
Cos*θ* = cosine of angle between y=x, and regression line, for each set of experimental and calculated ^3^J*_HN_*_−_*_Hα_*-Coupling values. a) A*β*42, b) Tau43, c) amylin and d) *α*S

**Table S1:** Cos*θ* = cosine of the angle between y=x, and regression line, for each set of experimental and calculated ^3^J*_HN_*_−_*_Hα_*-Coupling values.

| Force field water model combination | Aβ42 | | Tau43 | | Amylin | αS | |
| --- | --- | --- | --- | --- | --- | --- | --- |
|  | Expt. | Server | Expt. | Server | Server | Expt. | Server |
| C36-IDPSFF/TIP4P-Ew | 0.94 | 0.89 | 0.65 | 0.74 | 0.98 | 0.96 | 0.93 |
| A99SB-ILDN/TIP4P-Ew | 0.96 | 0.87 | 0.52 | 0.82 | 0.84 | 0.89 | 0.86 |
| A99SB-ILDN-Dyes/TIP4P-Ew | 0.97 | 0.92 | 0.59 | 0.88 | 0.98 | 0.88 | 0.84 |
| C36-IDPSFF/A99SB-disp | 0.94 | 0.93 | 0.68 | 0.74 | 0.99 | 0.96 | 0.95 |
| FF99SB/OPC | 0.96 | 0.88 | 0.61 | 0.86 | 0.97 | 0.93 | 0.90 |
| A99SB-disp/A99SB-disp | 0.92 | 0.89 | 0.74 | 0.77 | 0.96 | 0.88 | 0.84 |
| FF14SB/TIP3P | 0.87 | 0.80 | 0.70 | 0.87 | 0.91 | 0.82 | 0.81 |
| CHARMM27/TIP3P | 0.88 | 0.77 | 0.60 | 0.75 | 0.98 | 0.85 | 0.82 |
| GROMOS 96 (54a7)/SPC/E | 0.94 | 0.86 | 0.51 | 0.67 | 0.83 | 0.68 | 0.68 |
| CHARMM36m/CHARMM TIP3P | 0.92 | 0.84 | 0.63 | 0.64 | 0.94 | 0.55 | 0.54 |
| CHARMM36m/TIP3P | 0.93 | 0.87 | 0.56 | 0.48 | 0.94 | 0.81 | 0.75 |
| A14SB/TIP4P-Ew | 0.92 | 0.88 | 0.67 | 0.80 | 0.93 | 0.85 | 0.81 |
| CHARMM27/CHARMM TIP3P | 0.88 | 0.75 | 0.62 | 0.64 | 0.92 | 0.87 | 0.79 |

**Table S2:** Mean squared error (MSE) between Experimental/*RC_3JHNHa Server* and Calculated ^3^J*_HN_*_−_*_Hα_*-Coupling Constants of IDPs Using the Parameters of the Karplus Equations.

| Force field water model combination | Aβ42 | | Tau43 | | Amylin | αS | |
| --- | --- | --- | --- | --- | --- | --- | --- |
|  | Expt. | Server | Expt. | Server | Server | Expt. | Server |
| C36-IDPSFF/TIP4P-Ew | 0.69 | 0.50 | 2.70 | 0.43 | 0.35 | 0.32 | 0.38 |
| A99SB-ILDN/TIP4P-Ew | 0.63 | 0.64 | 2.53 | 0.59 | 0.35 | 0.68 | 0.63 |
| A99SB-ILDN-Dyes/TIP4P-Ew | 0.63 | 0.53 | 2.54 | 0.62 | 0.32 | 0.68 | 0.64 |
| C36-IDPSFF/A99SB-disp | 0.65 | 0.41 | 2.71 | 0.44 | 0.34 | 0.24 | 0.29 |
| FF99SB/OPC | 0.62 | 0.61 | 2.41 | 0.60 | 0.41 | 0.69 | 0.63 |
| A99SB-disp/A99SB-disp | 1.42 | 0.88 | 4.23 | 0.84 | 1.11 | 0.59 | 0.74 |
| FF14SB/TIP3P | 1.68 | 1.06 | 4.44 | 0.82 | 1.11 | 0.59 | 0.66 |
| CHARMM27/TIP3P | 1.43 | 0.94 | 3.47 | 0.61 | 0.71 | 0.51 | 0.59 |
| GROMOS 96 (54a7)/SPC/E | 1.62 | 0.83 | 4.78 | 0.86 | 0.92 | 0.91 | 0.93 |
| CHARMM36m/CHARMM TIP3P | 1.16 | 0.68 | 3.39 | 0.57 | 0.53 | 1.05 | 1.09 |
| CHARMM36m/TIP3P | 0.68 | 1.15 | 3.73 | 0.70 | 0.61 | 0.58 | 0.68 |
| A14SB/TIP4P-Ew | 1.63 | 0.95 | 4.18 | 0.72 | 0.96 | 0.58 | 0.71 |
| CHARMM27/CHARMM TIP3P | 1.48 | 0.97 | 3.27 | 0.65 | 0.64 | 0.45 | 0.55 |

**Table S3:** Mean absolute error (MAE) between Experimental/*RC_3JHNHa Server* and Calculated ^3^J*_HN_*_−_*_Hα_*-Coupling Constants of IDPs.

| Force field water model combination | Aβ42 | | Tau43 | | Amylin | αS | |
| --- | --- | --- | --- | --- | --- | --- | --- |
|  | Expt. | Server | Expt. | Server | Server | Expt. | Server |
| C36-IDPSFF/TIP4P-Ew | 0.70 | 0.48 | 1.36 | 0.42 | 0.43 | 0.44 | 0.45 |
| A99SB-ILDN/TIP4P-Ew | 0.69 | 0.58 | 1.20 | 0.59 | 0.43 | 0.67 | 0.63 |
| A99SB-ILDN-Dyes/TIP4P-Ew | 0.68 | 0.54 | 1.18 | 0.59 | 0.37 | 0.66 | 0.64 |
| C36-IDPSFF/A99SB-disp | 0.67 | 0.42 | 1.37 | 0.43 | 0.41 | 0.37 | 0.38 |
| FF99SB/OPC | 0.68 | 0.59 | 1.17 | 0.61 | 0.51 | 0.69 | 0.64 |
| A99SB-disp/A99SB-disp | 1.03 | 0.81 | 1.88 | 0.76 | 0.93 | 0.63 | 0.68 |
| FF14SB/TIP3P | 1.11 | 0.86 | 1.89 | 0.73 | 0.93 | 0.61 | 0.64 |
| CHARMM27/TIP3P | 1.01 | 0.81 | 1.58 | 0.60 | 0.69 | 0.56 | 0.61 |
| GROMOS 96 (54a7)/SPC/E | 1.10 | 0.75 | 1.88 | 0.71 | 0.81 | 0.79 | 0.78 |
| CHARMM36m/CHARMM TIP3P | 0.91 | 0.66 | 1.58 | 0.53 | 0.55 | 0.84 | 0.85 |
| CHARMM36m/TIP3P | 0.89 | 0.62 | 1.66 | 0.63 | 0.54 | 0.61 | 0.64 |
| A14SB/TIP4P-Ew | 1.12 | 0.85 | 1.84 | 0.71 | 0.85 | 0.62 | 0.69 |
| CHARMM27/CHARMM TIP3P | 1.00 | 0.74 | 1.52 | 0.61 | 0.66 | 0.55 | 0.57 |

### S4 ^3^J*_HN_*_−_*_Hα_*-Coupling values

**Table S4:**
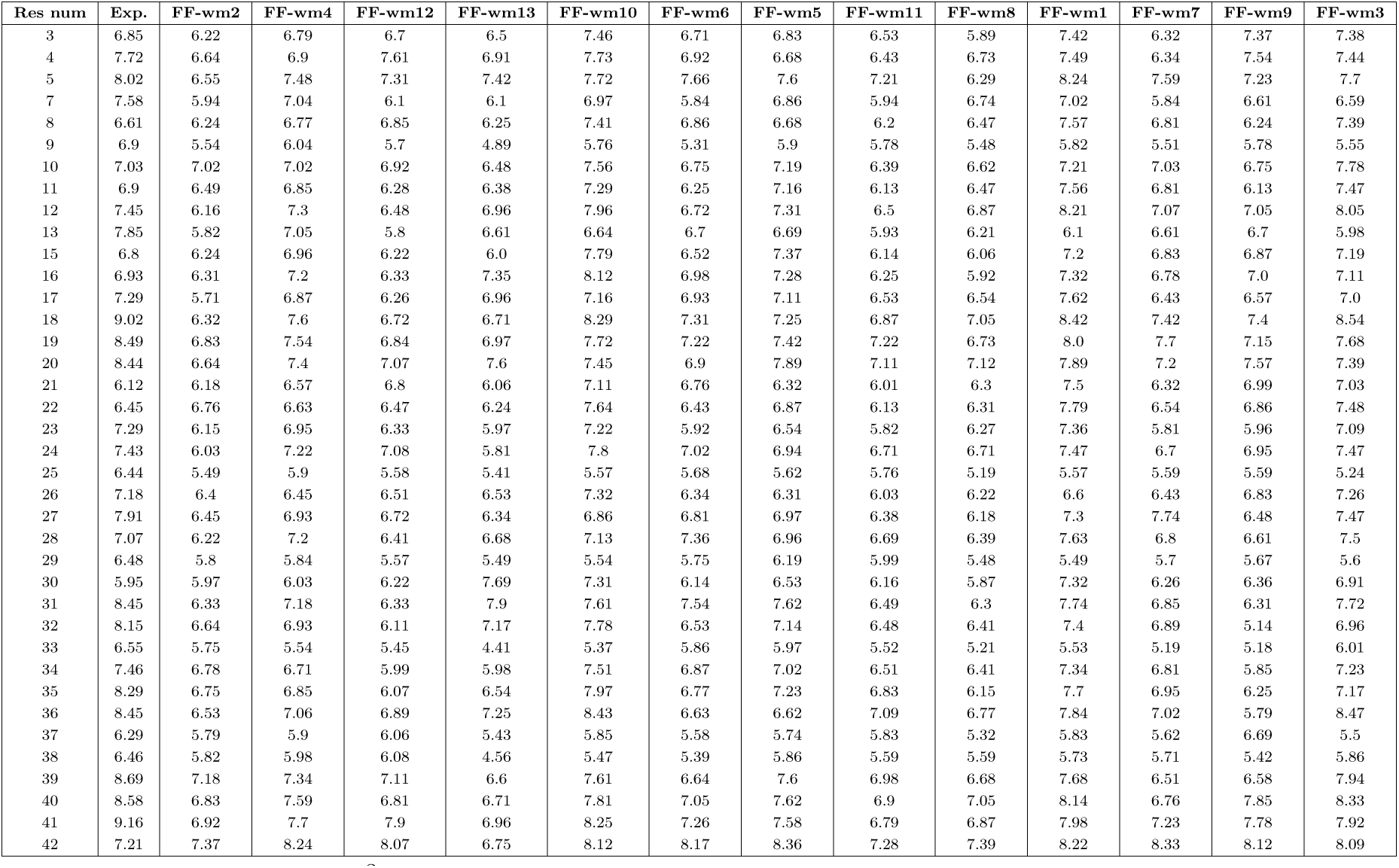
Experimental ^3^J*_HN_*_−_*_Hα_*-Coupling Values from [63] and corresponding Calculated data for A*β*42.

**Table S5:**
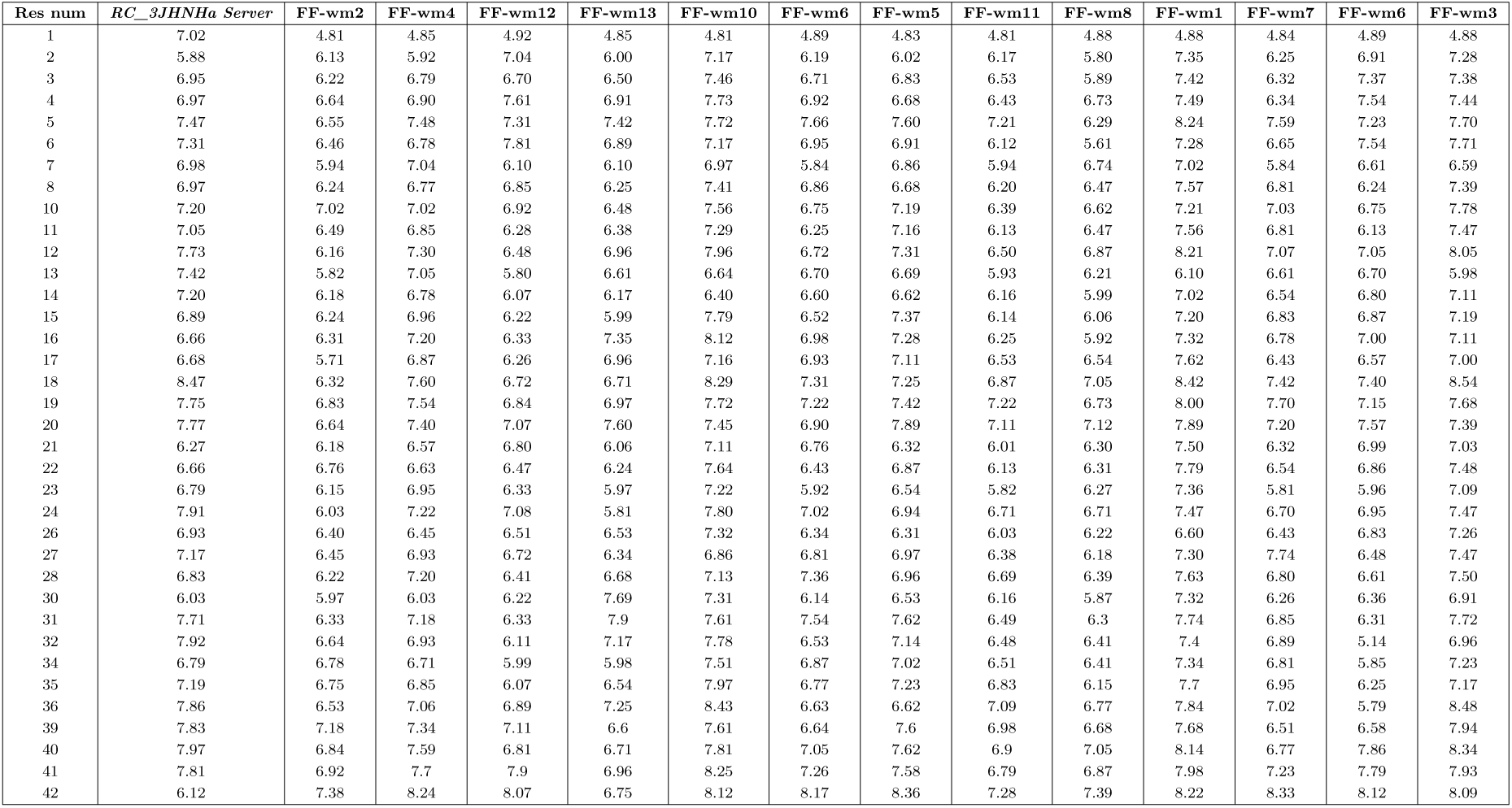
Predicted (*RC_3JHNHa Server*) ^3^J*_HN_*_−_*_Hα_*-Coupling Values [64] and corresponding Calculated data for A*β*42.

**Table S6:**
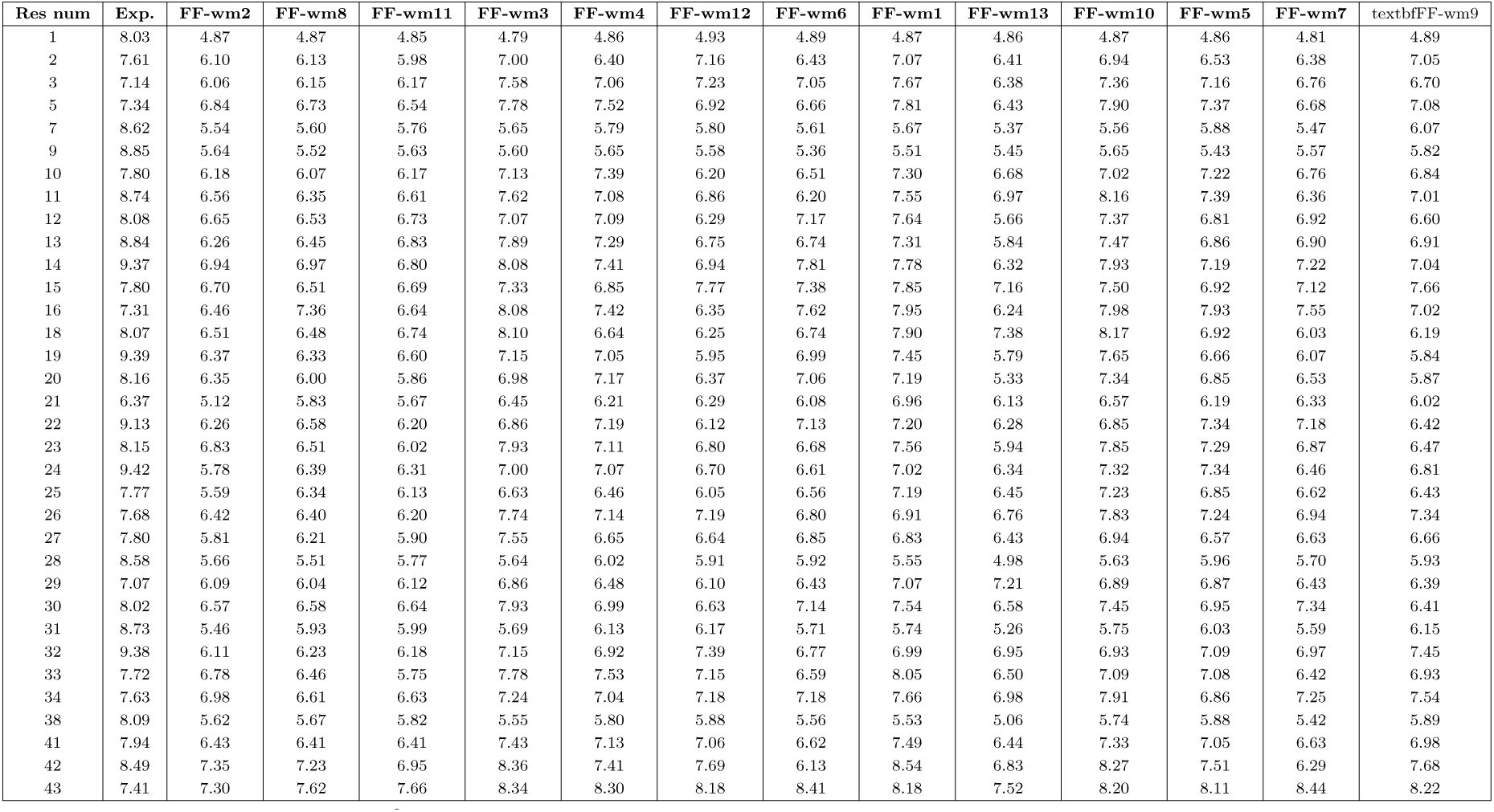
Experimental ^3^J*_HN_*_−_*_Hα_*-Coupling Values from [43] and corresponding Calculated data for Tau43.

**Table S7:**
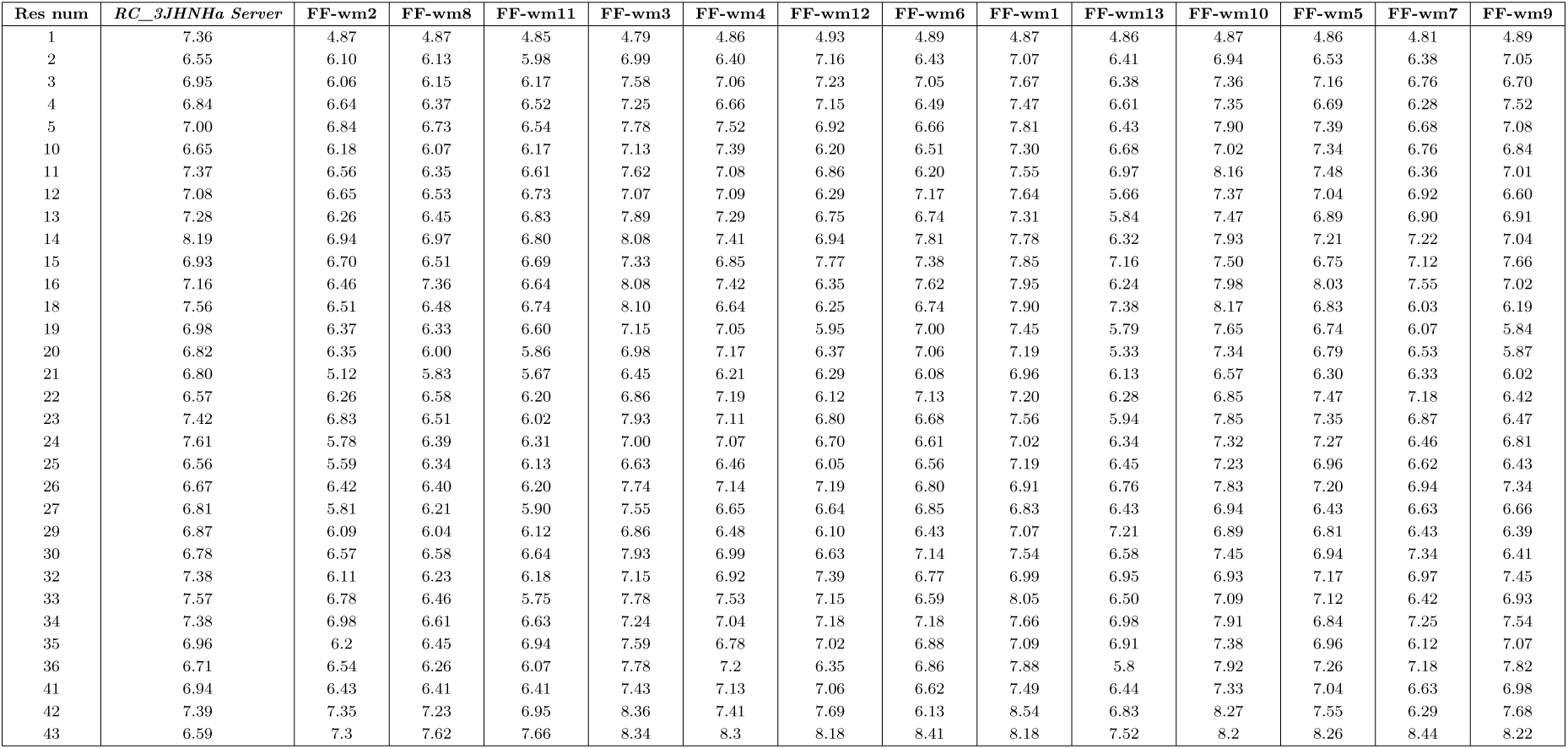
Predicted (*RC_3JHNHa Server*) ^3^J*_HN_*_−_*_Hα_*-Coupling Values [64] and corresponding Calculated data for Tau43.

**Table S8:**
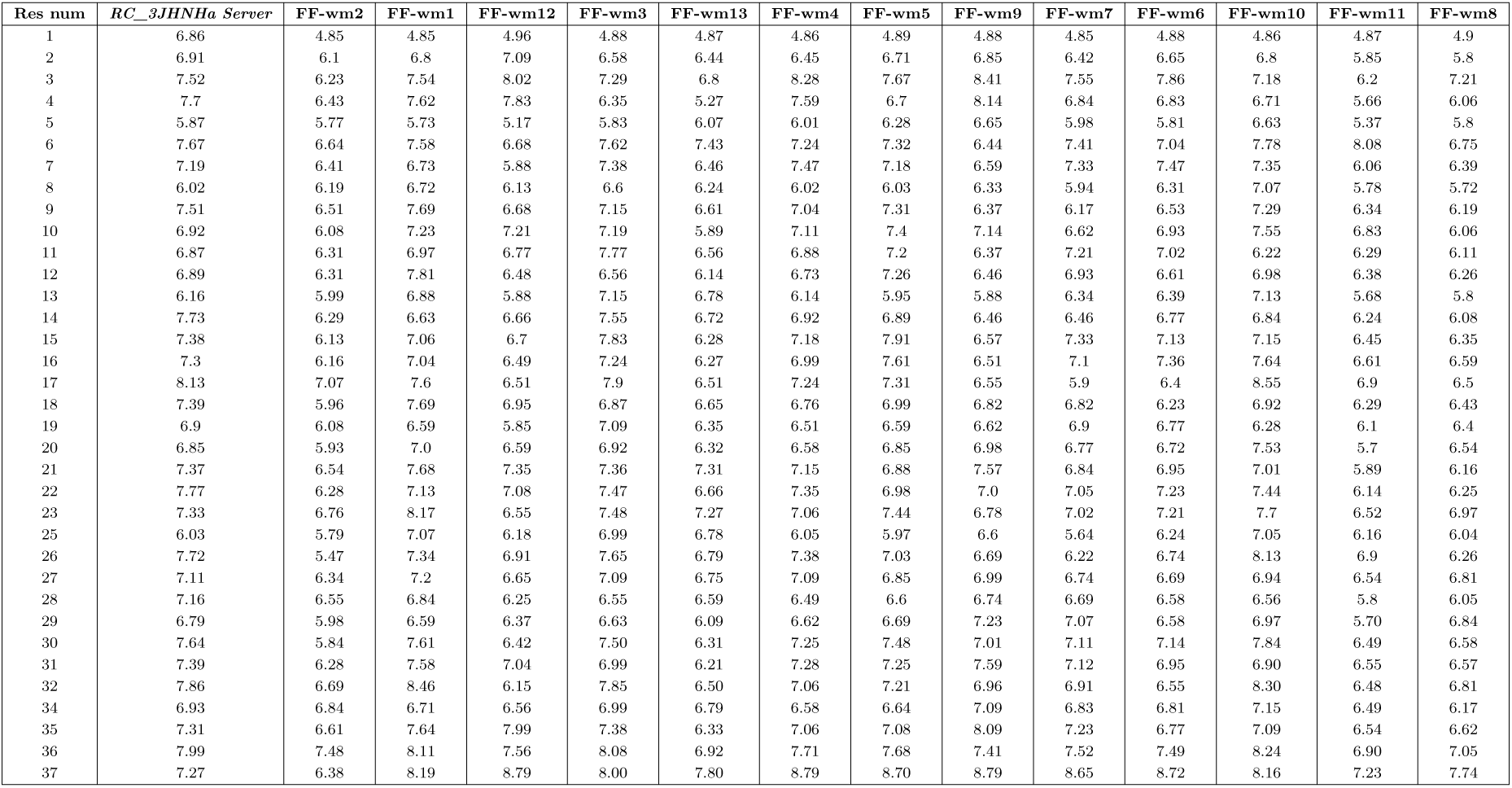
Predicted (*RC_3JHNHa Server*) ^3^J*_HN_*_−_*_Hα_*-Coupling Values [64] and corresponding Calculated data for Amylin.

**Table S9:**
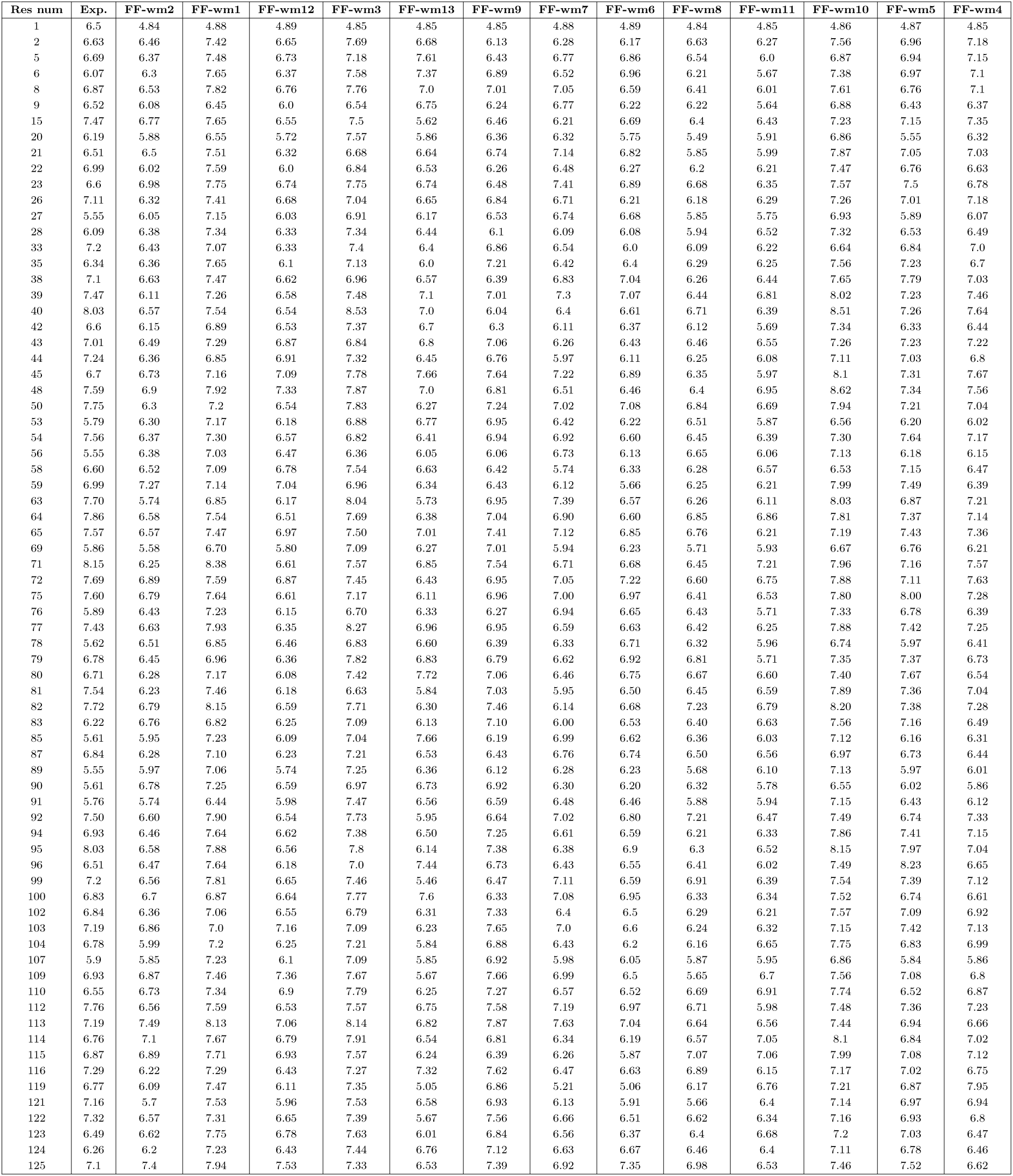
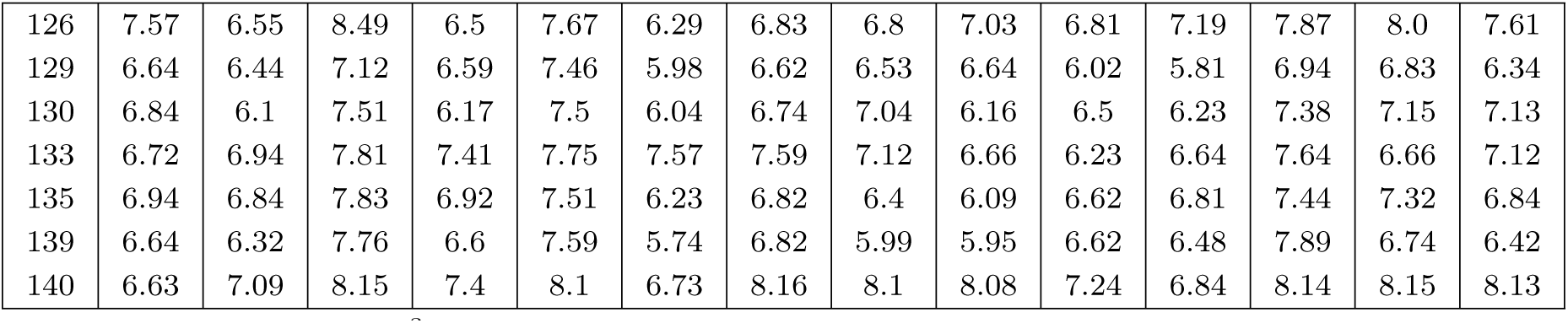
Experimental ^3^J*_HN_*_−_*_Hα_*-Coupling Values from [62] and corresponding Calculated data for *α*S.

**Table S10:**
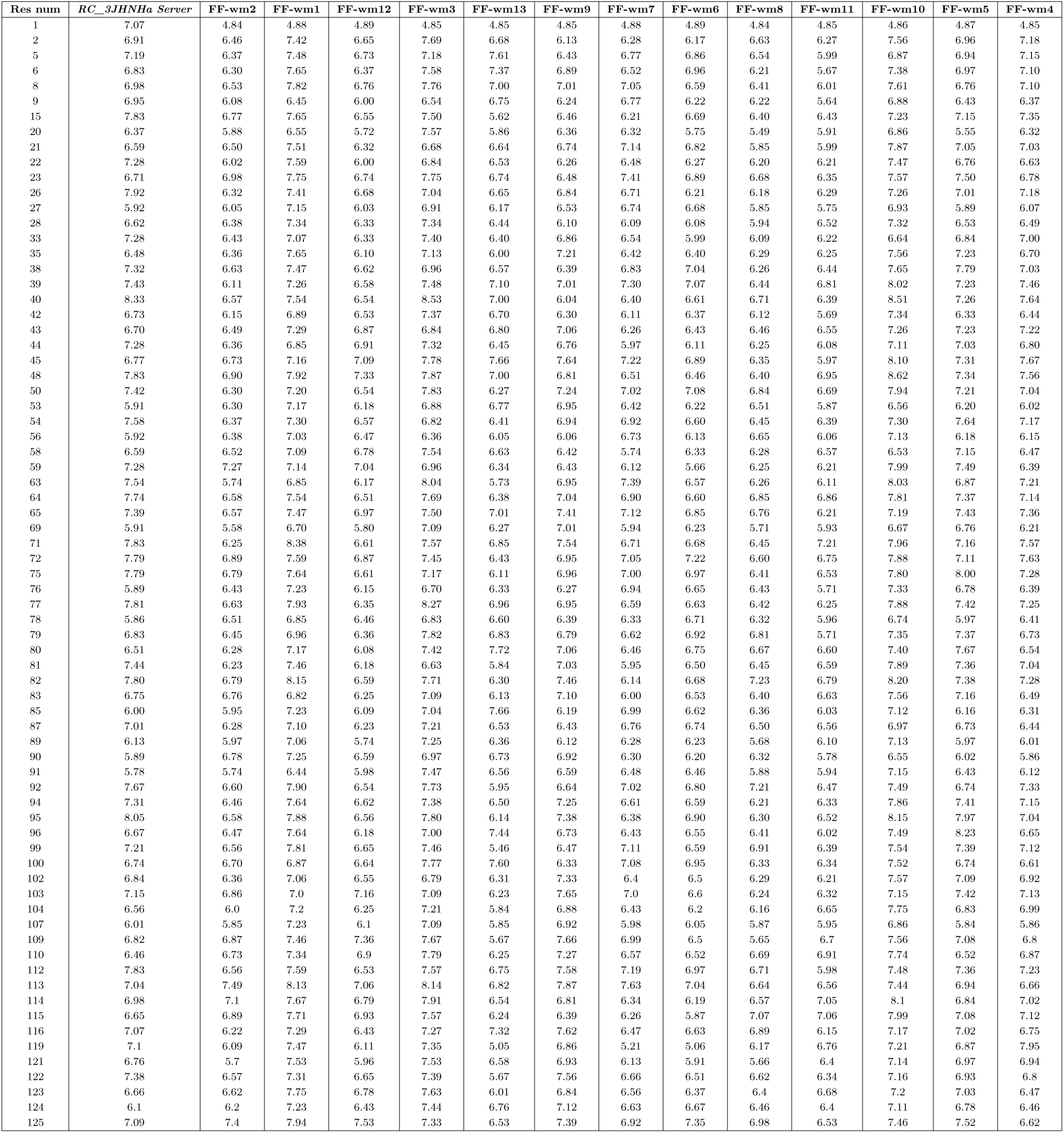
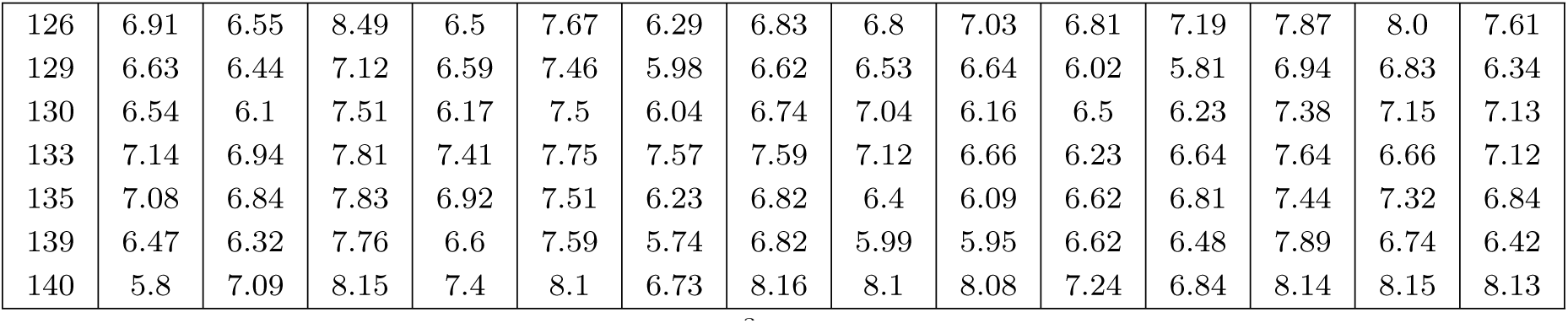
Predicted (*RC_3JHNHa Server*) ^3^J*_HN_*_−_*_Hα_*-Coupling Values [64] and corresponding Calculated data for *α*S.

### S5 2D Density Plots

**Fig. S4:**
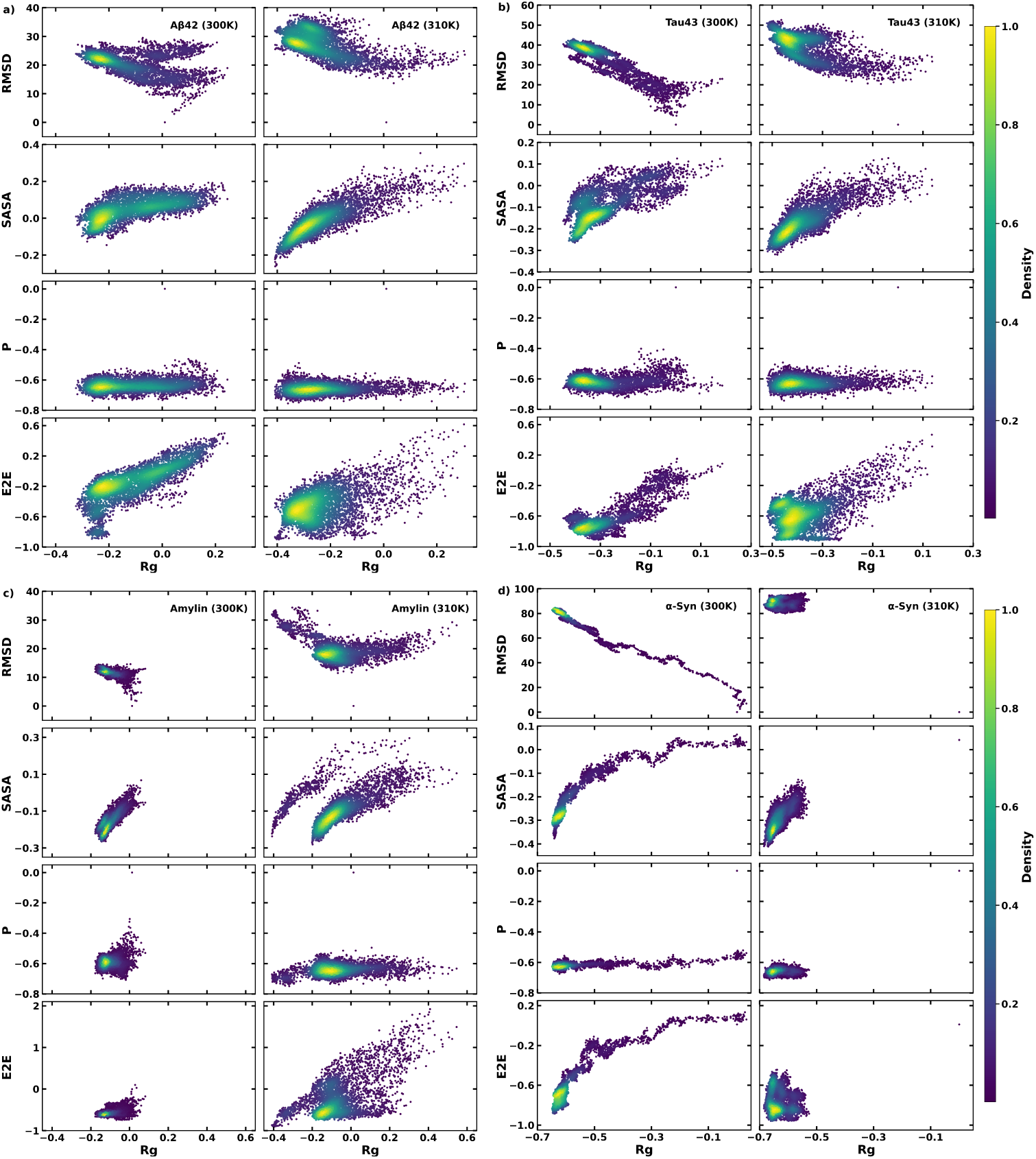
The 2D Kernel density plots as a function of Rg vs RMSD, Rg vs SASA, Rg vs P and Rg vs E2E as the coordinates for four selected proteins: a) A*β*42, b) Tau43, c) amylin and d) *α*S, for FF-wm1.

**Fig. S5:**
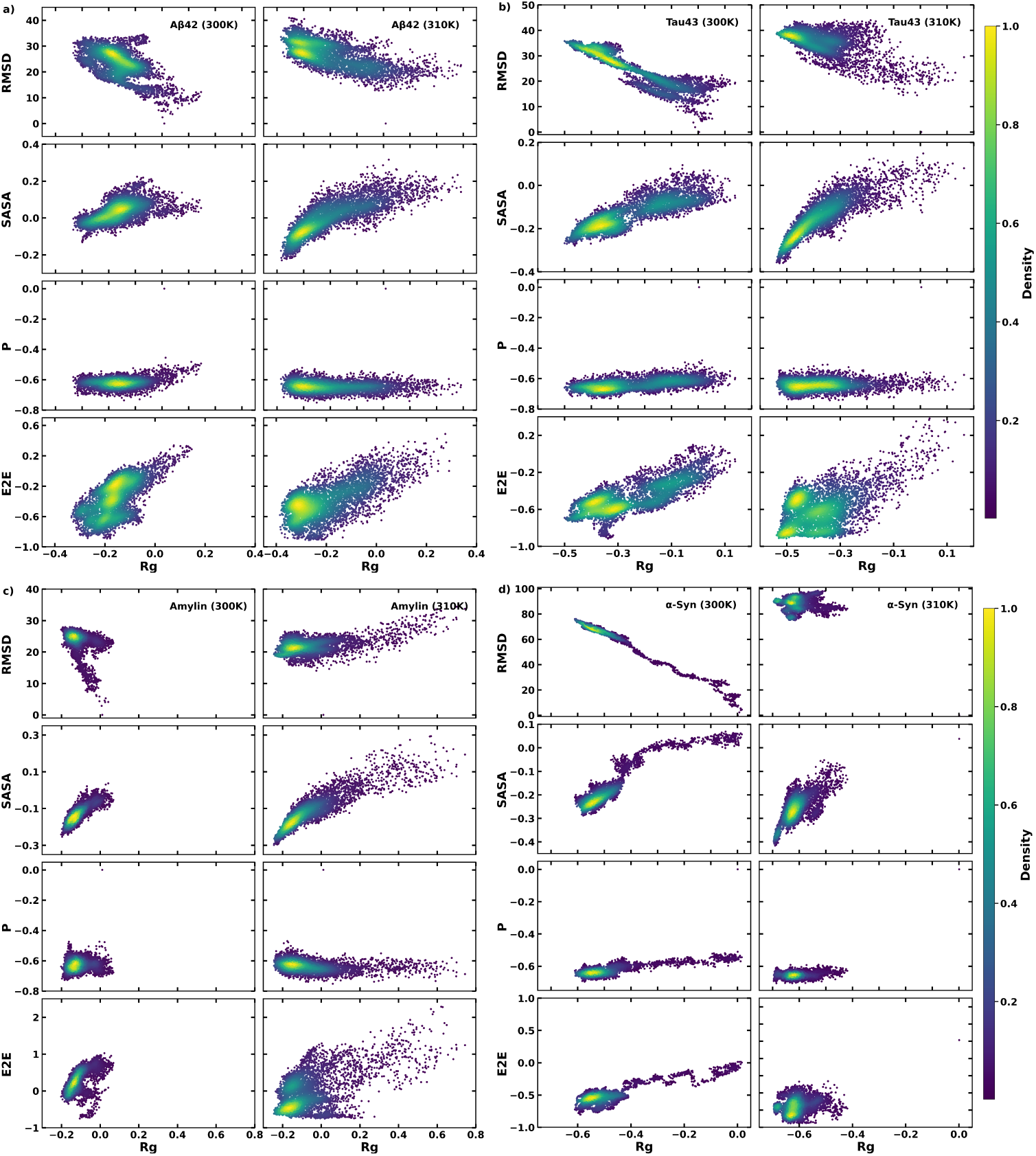
The 2D Kernel density plots as a function of Rg vs RMSD, Rg vs SASA, Rg vs P and Rg vs E2E as the coordinates for four selected proteins: a) A*β*42, b) Tau43, c) amylin and d) *α*S, for FF-wm3.

**Fig. S6:**
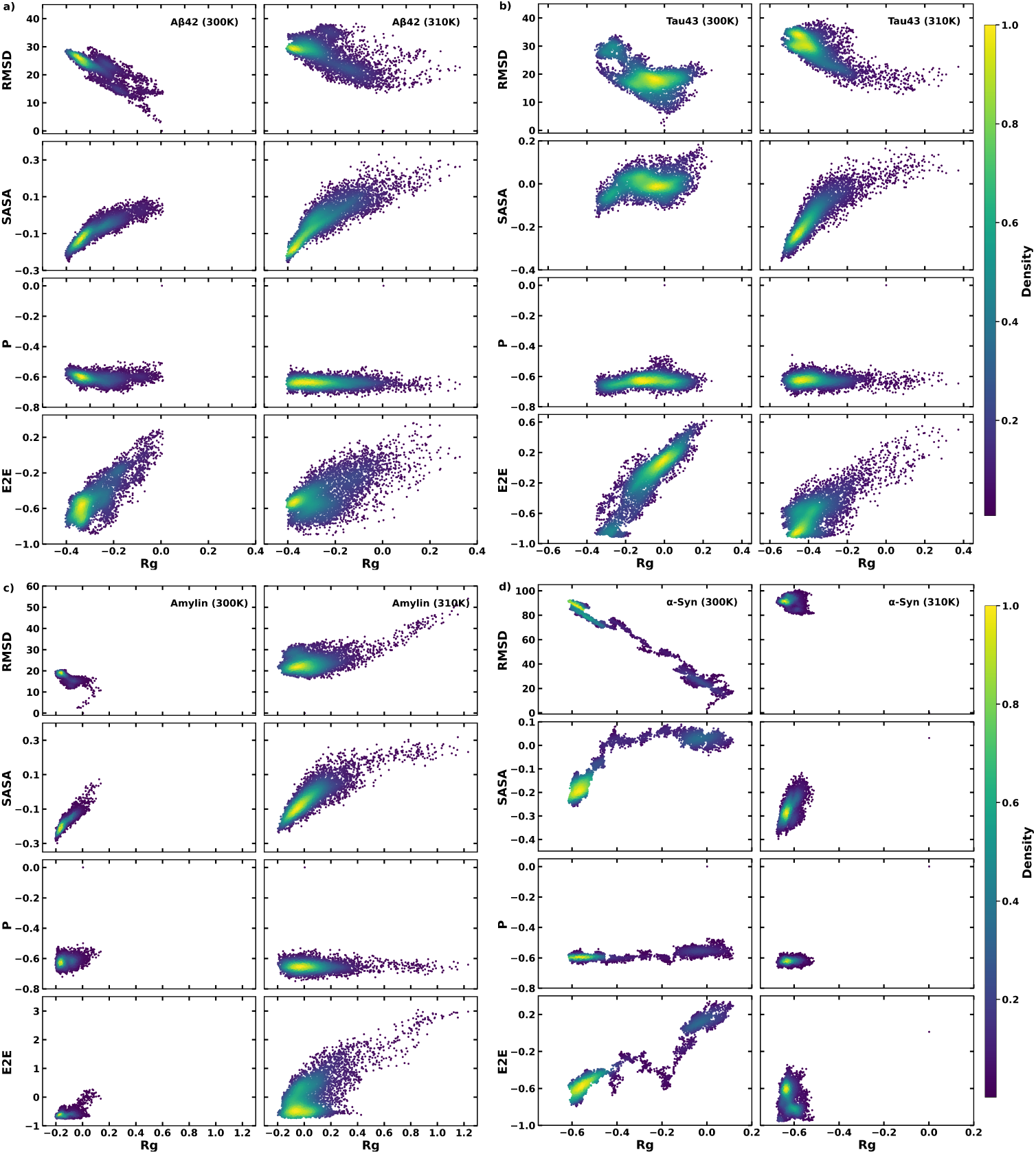
The 2D Kernel density plots as a function of Rg vs RMSD, Rg vs SASA, Rg vs P and Rg vs E2E as the coordinates for four selected proteins: a) A*β*42, b) Tau43, c) amylin and d) *α*S), for FF-wm5.

**Table S11:** Diffusion coefficient (×10^−4^Å^2^*/*ps) of water molecules around proteins (Sh_1_). The table reports the average diffusion coefficients (from every 2 ps time-window, from 2-20 ps MSD data) and their standard deviations in round brackets for FF-wm5.

| Temp. | Aβ42 | Tau43 | Amylin | αS | HP36 | WW |
| --- | --- | --- | --- | --- | --- | --- |
| 300 K | $10.1 \pm 1.7$ | $11.3 \pm 1.6$ | $10.3 \pm 1.8$ | $9.12 \pm 1.5$ | $9.65 \pm 1.7$ | $9.89 \pm 1.7$ |
| 310 K | $12.4 \pm 1.6$ | $12.3 \pm 1.7$ | $12.3 \pm 1.7$ | $10.5 \pm 1.7$ | $11.4 \pm 1.8$ | $11.9 \pm 1.8$ |

**Table S12:** Diffusion coefficient (×10^−4^Å^2^*/*ps) of water molecules around proteins (Sh_1_). The table reports the average diffusion coefficients (from every 2 ps time-window, from 2-20 ps MSD data) and their standard deviations in round brackets for FF-wm1.

| Temp. | Aβ42 | Tau43 | Amylin | αS | HP36 | WW |
| --- | --- | --- | --- | --- | --- | --- |
| 300 K | $9.29 \pm 1.7$ | $10.5 \pm 1.6$ | $9.03 \pm 1.6$ | $8.87 \pm 1.5$ | $13.2 \pm 0.5$ | $9.55 \pm 1.7$ |
| 310 K | $11.6 \pm 1.8$ | $11.7 \pm 1.7$ | $10.9 \pm 1.8$ | $9.54 \pm 1.9$ | $11.6 \pm 1.8$ | $11.8 \pm 1.8$ |

**Table S13:** Diffusion coefficient (×10^−4^Å^2^*/*ps) of water molecules around proteins (Sh_1_). The table reports the average diffusion coefficients (from every 2 ps time window, from 2-20 ps MSD data) and their standard deviations in round brackets for FF-wm3.

| Temp. | Aβ42 | Tau43 | Amylin | αS | HP36 | WW |
| --- | --- | --- | --- | --- | --- | --- |
| 300 K | $9.10 \pm 1.6$ | $10.2 \pm 1.6$ | $10.4 \pm 1.7$ | $8.84 \pm 1.5$ | $9.86 \pm 1.7$ | $10.2 \pm 1.6$ |
| 310 K | $11.9 \pm 1.6$ | $12.5 \pm 1.7$ | $11.0 \pm 1.8$ | $10.1 \pm 1.7$ | $11.2 \pm 1.7$ | $12.0 \pm 1.8$ |

### S6 Mean Square Displacement (MSD)

**Fig. S7:**
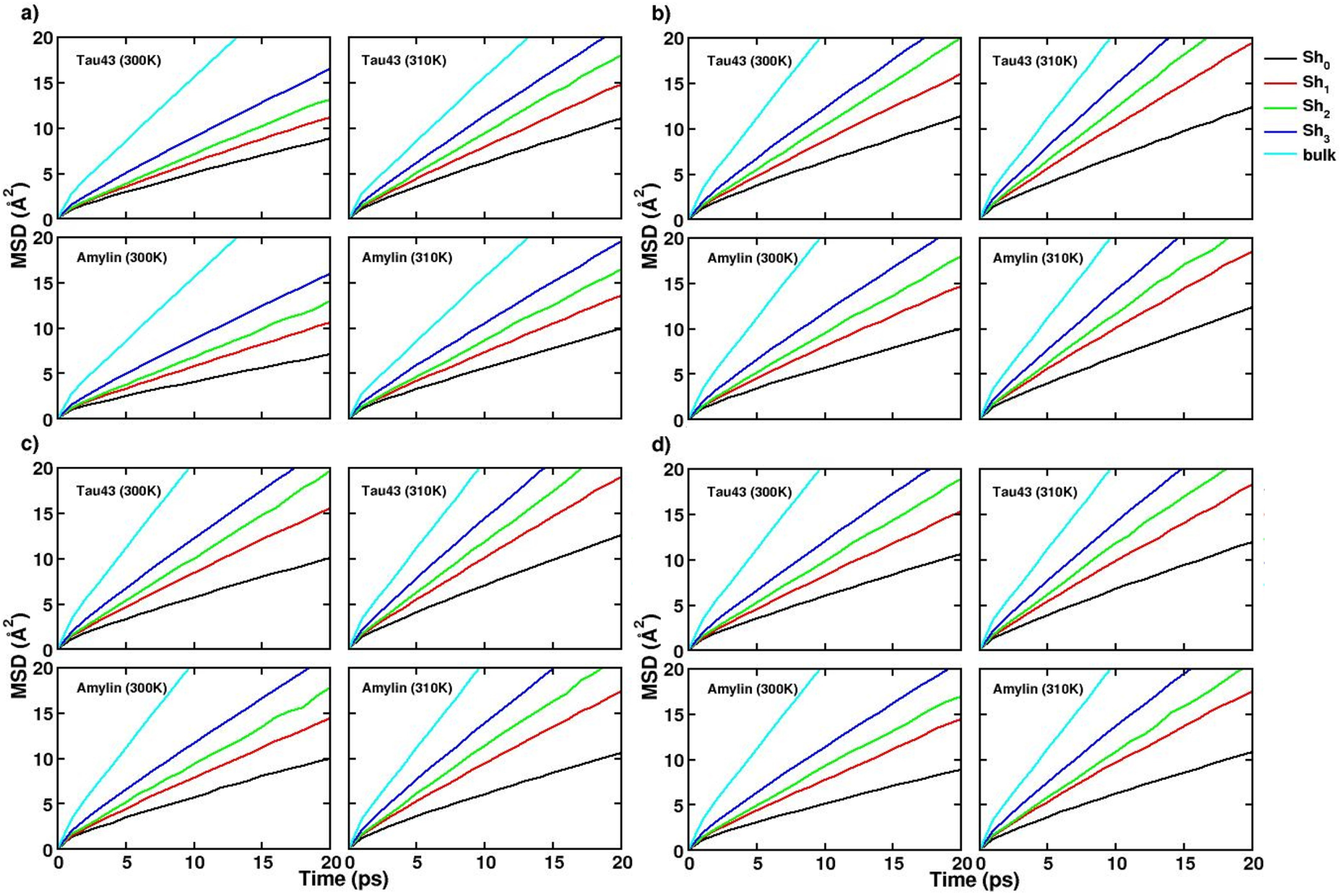
Mean squared displacement (MSD) of the hydration shells (Sh_1_, Sh_2_, Sh_3_, and Sh_4_) surrounding the proteins (Tau43 and Amylin) at temperature 300 K and 310 K, based on the hydration layer peaks identified using g(r) (see Fig.3) for a)FF-wm4, b)FF-wm5, c)FF-wm3 and d)FF-wm1.

**Fig. S8:**
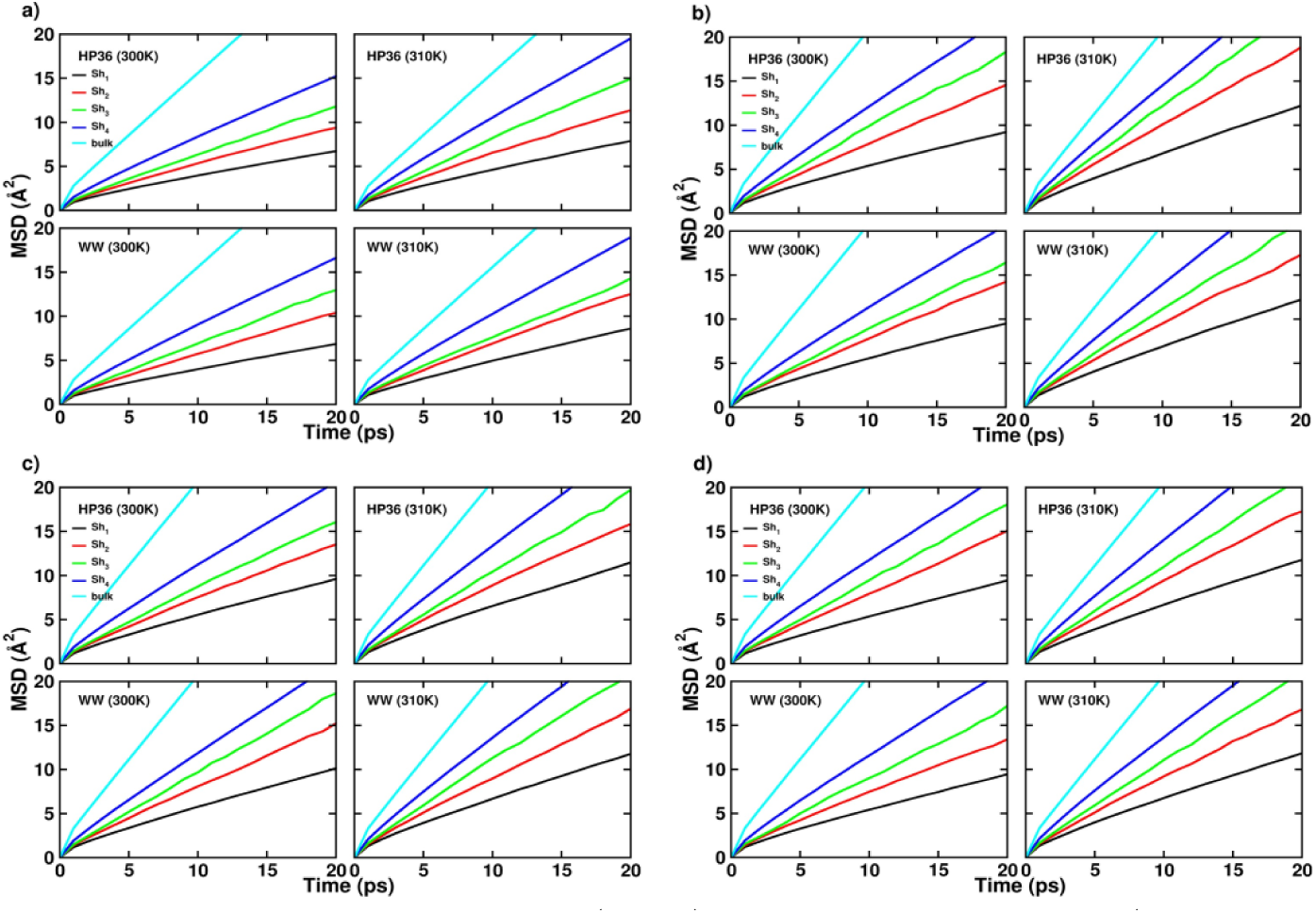
Mean squared displacement (MSD) of the hydration shells (Sh_1_, Sh_2_, Sh_3_, and Sh_4_) surrounding the proteins (HP36 and ww-domain) at temperature 300 K and 310 K, based on the hydration layer peaks identified using g(r) (see Fig.3) for a)FF-wm4, b)FF-wm5, c)FF-wm3 and d)FF-wm1.

#### S6.1 Diffusion Coefficient differences

D1_diff_ = MSD at 310 K - MSD at 300 K, for each shell (Sh_1_, Sh_2_, Sh_3_, Sh_4_) (Fig 5, Fig. S9),

D2_diff_ = MSD difference corresponding to each hydration shell at each temperature (300 K and 310 K).

There are 6 different D2_diffs_: Sh_2_-Sh_1_, Sh_3_-Sh_1_, Sh_3_-Sh_2_, Sh_4_-Sh_1_, Sh_4_-Sh_2_, Sh_4_-Sh_3_ corresponding to each temperature (Fig6, FigS11, FigS12).

**Fig. S9:**
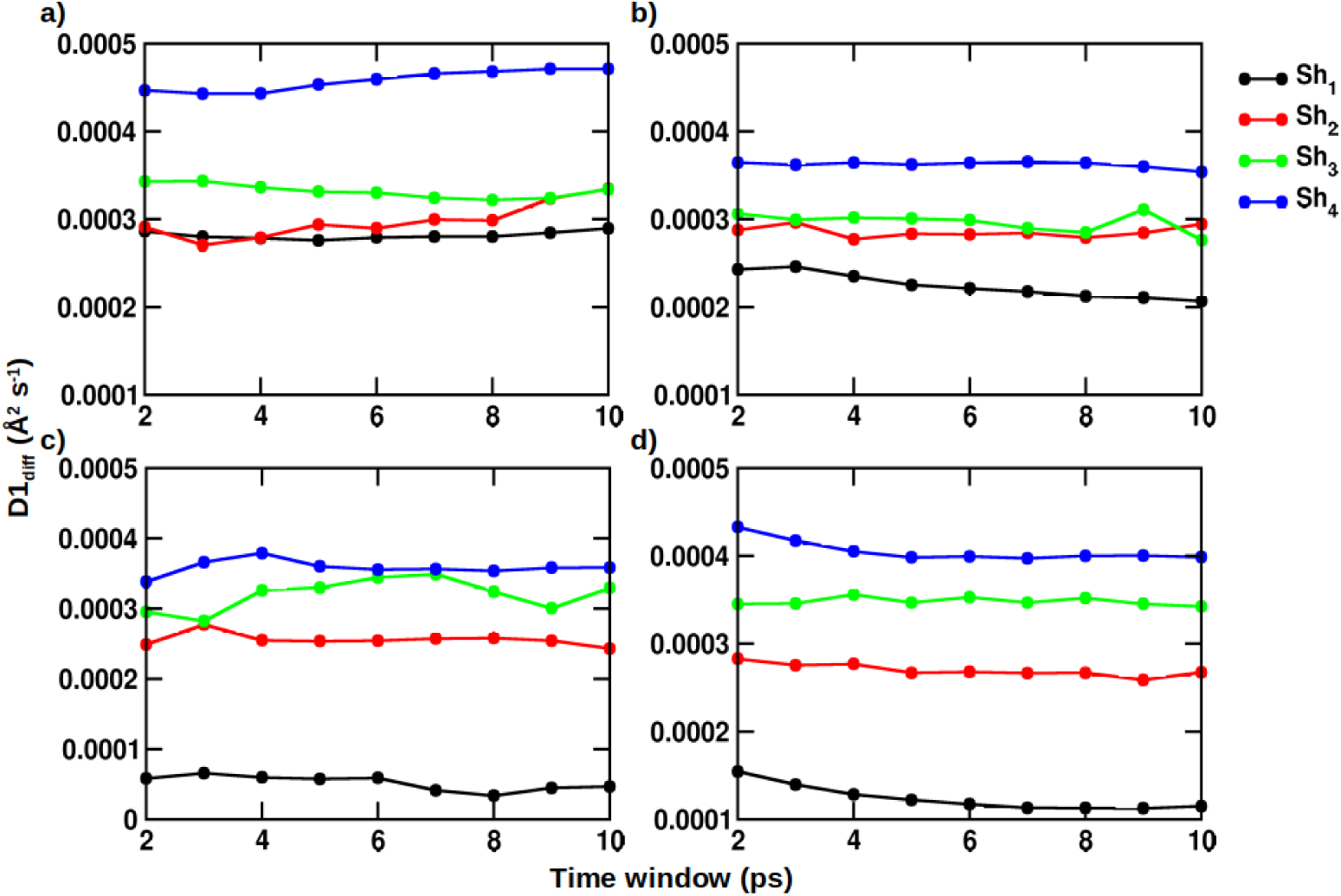
Differences in the mean squared displacement (D1_diff_) between the two simulated temperatures, for the respective hydration shells (see Fig. 3), tabulated at different time window, for a) A*β*42, b) Tau43, c) Amylin, d) *α*S, for FF-wm3 (see Table 1 for FF-wm combinations).

**Fig. S10:**
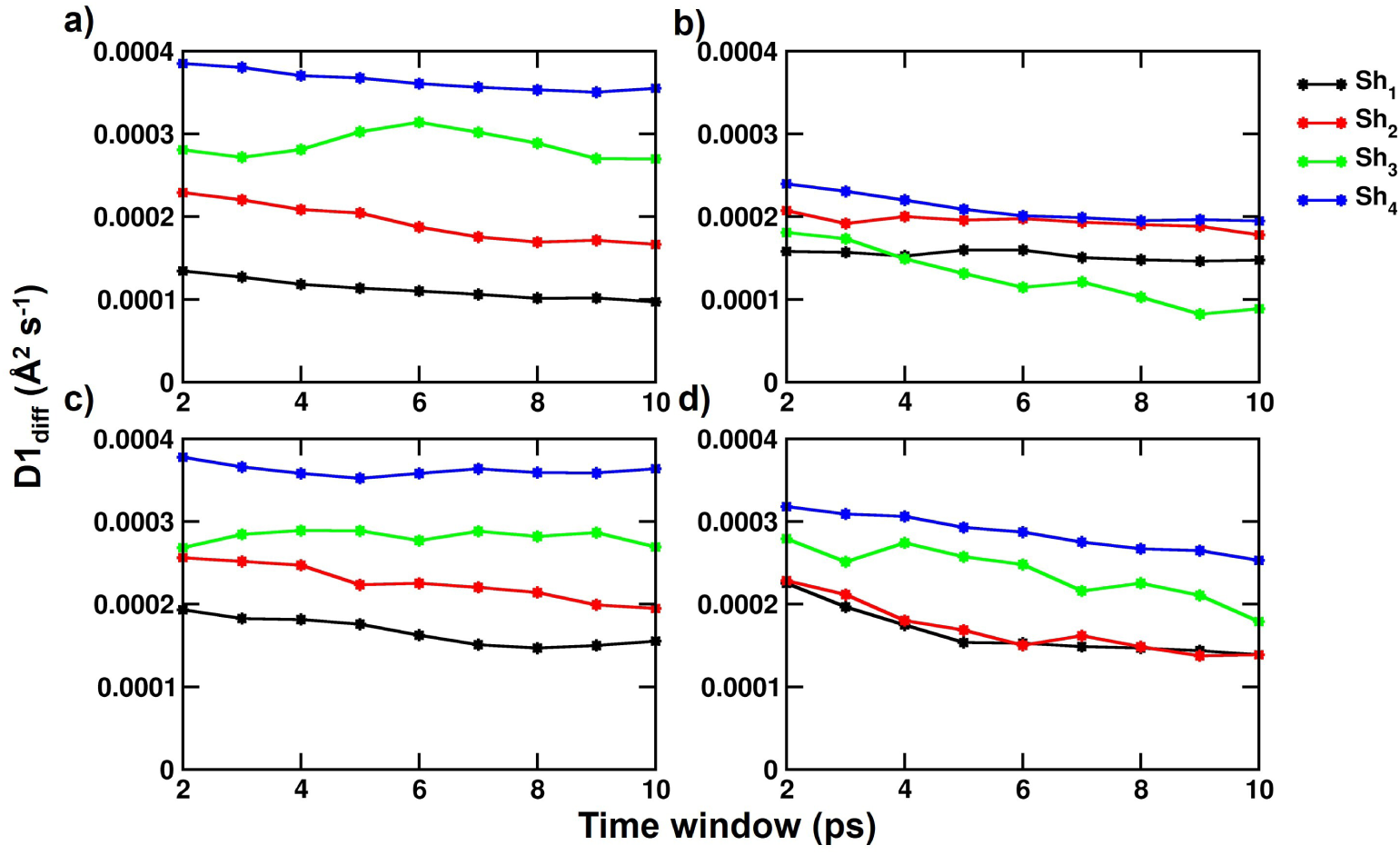
Differences in the mean squared displacement (D1_diff_) between the two simulated temperatures, for the respective hydration shells (see Fig. 3), tabulated at different time window (Table 3) for a) HP-36 with FF-wm4, b) WW-domain with FF-wm4, c) HP-36 with FF-wm5, d) WW-domain with FF-wm5 (see Table 1 for FF-wm combinations).

**Fig. S11:**
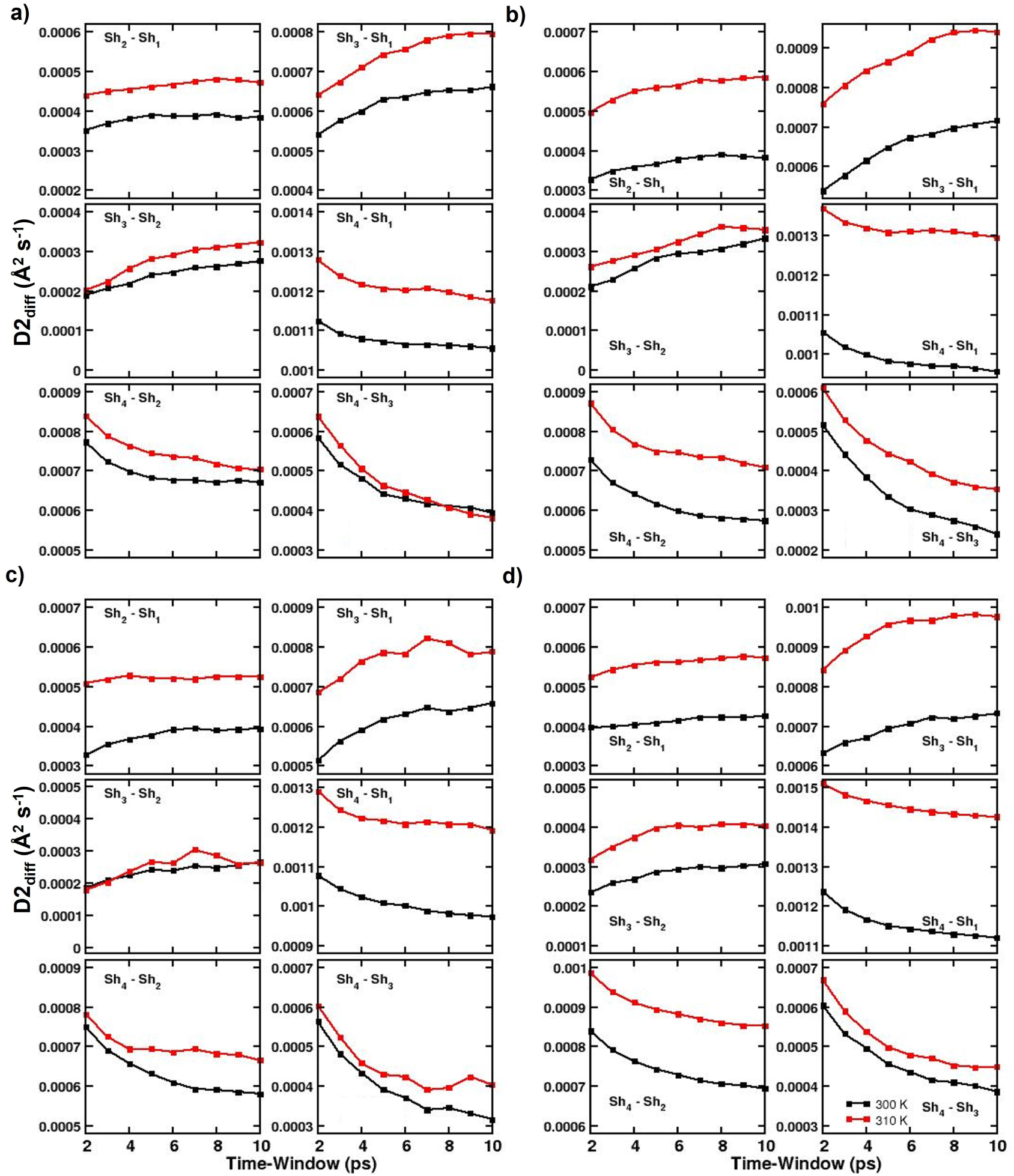
Differences in the mean squared displacement (D2_diff_) between the consecutive hydration shells (Fig.3), at the different time domains (see Table 3), for the two simulated temperatures, for a) A*β*42, b) Tau43, c) Amylin, d) *α*S, for FF-wm3 (see Table 1 for FF-wm combinations).

**Fig. S12:**
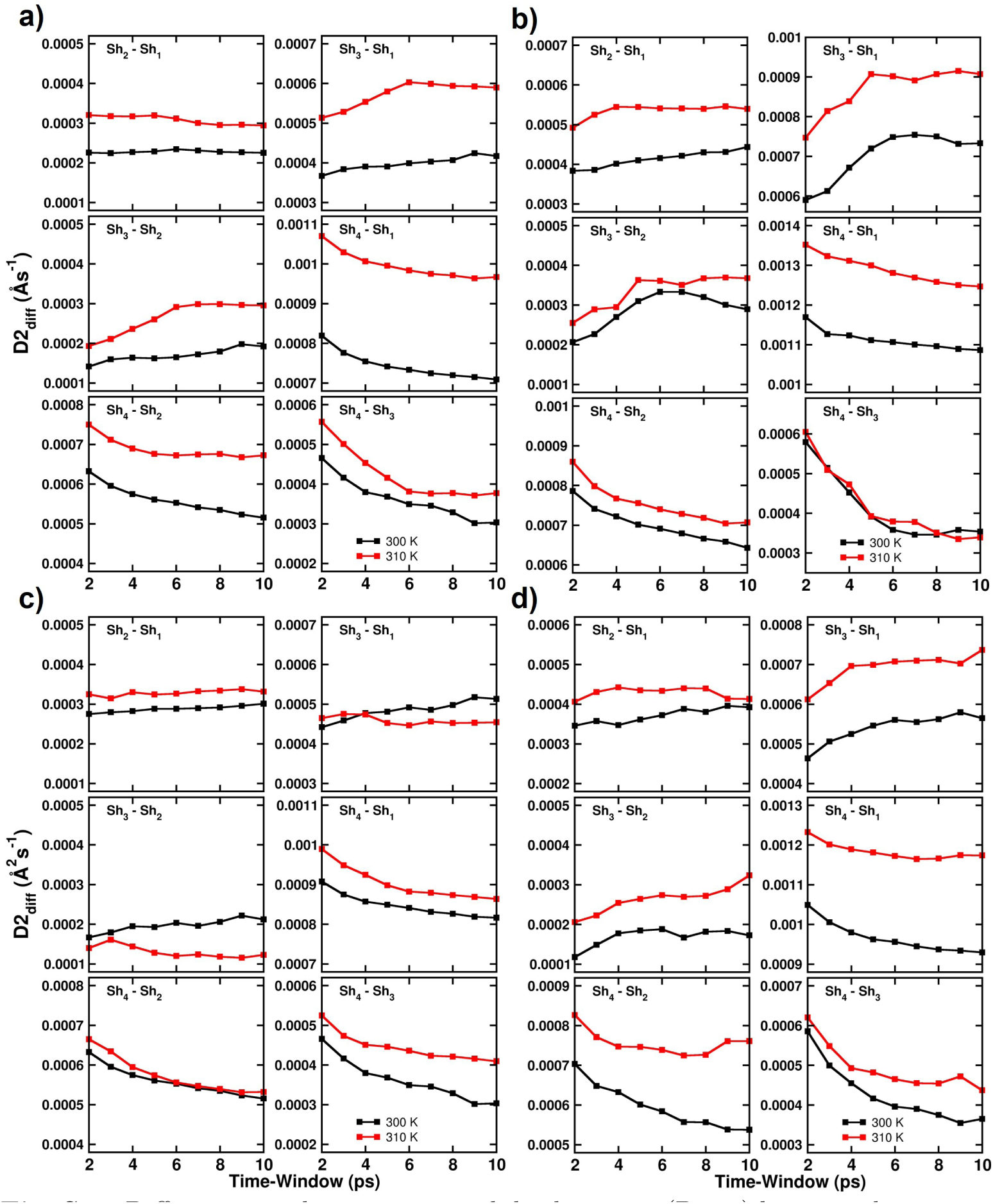
Differences in the mean squared displacement (D2_diff_) between the consecutive hydration shells (see Fig. 3), at different temperatures, tabulated at different time domains (Table 3), for a) HP-36 with FF-wm4. b) HP-36 with FF-wm3, c) WW-domain with FF-wm4, d) WW-domain with FF-wm3 (see Table 1 for FF-wm combinations).

**Table S14:** Diffusion Data (×10^−4^Å^2^*/*ps), for FF-wm4 using linear regression fitting over the MSD data (from 2-20 ps averaged over every 2 ps time-window)

| **Aβ42** | 300 K | 310 K | 300 K | 310 K | 300 K | 310 K | 300 K | 310 K |
| --- | --- | --- | --- | --- | --- | --- | --- | --- |
| **Shell** | **1–3 $\text{\AA}$** |  | **3–4 $\text{\AA}$** |  | **4–5 $\text{\AA}$** |  | **5–8 $\text{\AA}$** |  |
| 2 | 11.2 | 12.5 | 13.6 | 15.4 | 15.1 | 17.0 | 19.6 | 22.2 |
| 3 | 9.31 | 10.6 | 11.7 | 13.5 | 13.3 | 15.3 | 17.3 | 19.8 |
| 4 | 8.42 | 9.68 | 10.9 | 12.6 | 12.6 | 14.4 | 16.3 | 18.6 |
| 5 | 7.86 | 9.10 | 10.3 | 12.1 | 12.1 | 14.1 | 15.6 | 17.9 |
| 6 | 7.49 | 8.72 | 10.0 | 11.7 | 11.9 | 13.8 | 15.2 | 17.5 |
| 7 | 7.21 | 8.42 | 9.74 | 11.4 | 11.7 | 13.6 | 14.8 | 17.1 |
| 8 | 6.99 | 8.15 | 9.55 | 11.2 | 11.7 | 13.4 | 14.6 | 16.8 |
| 9 | 6.84 | 7.94 | 9.44 | 11.0 | 11.6 | 13.2 | 14.4 | 16.6 |
| 10 | 6.72 | 7.77 | 9.33 | 10.9 | 11.5 | 13.0 | 14.2 | 16.4 |
| **Average** | 8.00 | 9.20 | 10.5 | 12.2 | 12.4 | 14.2 | 15.8 | 18.1 |
| **Std-dev** | $\pm 1.4$ | $\pm 1.5$ | $\pm 1.4$ | $\pm 1.4$ | $\pm 1.1$ | $\pm 1.2$ | $\pm 1.7$ | $\pm 1.8$ |
| **Amylin** | 300 K | 310 K | 300 K | 310 K | 300 K | 310 K | 300 K | 310 K |
| **Shell** | **1–3 $\text{\AA}$** |  | **3–4 $\text{\AA}$** |  | **4–5 $\text{\AA}$** |  | **5–8 $\text{\AA}$** |  |
| 2 | 10.6 | 13.2 | 13.3 | 16.1 | 14.7 | 17.5 | 19.1 | 22.4 |
| 3 | 8.60 | 11.2 | 11.3 | 14.1 | 12.9 | 15.7 | 16.8 | 19.9 |
| 4 | 7.68 | 10.2 | 10.4 | 13.1 | 12.0 | 14.8 | 15.6 | 18.7 |
| 5 | 7.11 | 9.60 | 9.90 | 12.5 | 1.16 | 14.5 | 14.9 | 17.9 |
| 6 | 6.76 | 9.22 | 9.61 | 12.2 | 11.4 | 14.3 | 14.5 | 17.5 |
| 7 | 6.49 | 8.94 | 9.33 | 11.9 | 11.1 | 14.3 | 14.1 | 17.1 |
| 8 | 6.30 | 8.72 | 9.17 | 11.7 | 10.9 | 14.0 | 13.9 | 16.8 |
| 9 | 6.14 | 8.55 | 9.02 | 11.6 | 11.0 | 13.9 | 13.6 | 16.6 |
| 10 | 6.00 | 8.39 | 8.93 | 11.4 | 10.7 | 13.8 | 13.4 | 16.4 |
| **Average** | 7.29 | 9.77 | 10.1 | 12.7 | 11.8 | 14.8 | 15.1 | 18.1 |
| **Std-dev** | $\pm 1.4$ | $\pm 1.5$ | $\pm 1.4$ | $\pm 1.5$ | $\pm 1.2$ | $\pm 1.1$ | $\pm 1.8$ | $\pm 1.9$ |
| **tau43** | 300 K | 310 K | 300 K | 310 K | 300 K | 310 K | 300 K | 310 K |
| **Shell** | **1–3 $\text{\AA}$** |  | **3–4 $\text{\AA}$** |  | **4–5 $\text{\AA}$** |  | **5–8 $\text{\AA}$** |  |
| 2 | 11.9 | 14.0 | 14.0 | 16.8 | 14.8 | 18.5 | 19.3 | 23.2 |
| 3 | 10.0 | 12.0 | 12.1 | 15.0 | 13.1 | 16.9 | 17.0 | 20.9 |
| 4 | 9.13 | 11.1 | 11.2 | 14.1 | 12.5 | 16.2 | 15.9 | 19.8 |
| 5 | 8.67 | 10.6 | 10.7 | 13.6 | 12.1 | 15.9 | 15.3 | 19.2 |
| 6 | 8.32 | 10.2 | 10.3 | 13.1 | 11.8 | 15.6 | 14.8 | 18.8 |
| 7 | 8.05 | 9.92 | 10.0 | 12.9 | 11.6 | 15.5 | 14.5 | 18.4 |
| 8 | 7.82 | 9.66 | 9.83 | 12.7 | 11.4 | 15.5 | 14.2 | 18.2 |
| 9 | 7.64 | 9.46 | 9.60 | 12.6 | 11.3 | 15.2 | 14.0 | 18.0 |
| 10 | 7.47 | 9.30 | 9.47 | 12.4 | 11.2 | 15.1 | 13.8 | 17.8 |
| **Average** | 8.79 | 10.7 | 10.8 | 13.7 | 12.2 | 16.1 | 15.4 | 19.4 |
| **Std-dev** | $\pm 1.4$ | $\pm 1.5$ | $\pm 1.4$ | $\pm 1.4$ | $\pm 1.1$ | $\pm 1.0$ | $\pm 1.7$ | $\pm 1.7$ |
| $\alpha\mathbf{S}$ | 300 K | 310 K | 300 K | 310 K | 300 K | 310 K | 300 K | 310 K |
| **Shell** | **1–3 Å** |  | **3–4 Å** |  | **4–5 Å** |  | **5–8 Å** |  |
| 2 | 10.7 | 11.8 | 13.2 | 15.1 | 14.8 | 17.3 | 19.5 | 22.7 |
| 3 | 8.83 | 9.87 | 11.4 | 13.2 | 13.1 | 15.7 | 17.3 | 20.3 |
| 4 | 7.99 | 8.95 | 10.6 | 12.4 | 12.4 | 14.9 | 16.2 | 19.2 |
| 5 | 7.49 | 8.39 | 10.1 | 11.9 | 12.0 | 14.4 | 15.6 | 18.6 |
| 6 | 7.15 | 8.02 | 9.81 | 11.6 | 11.8 | 14.2 | 15.2 | 18.1 |
| 7 | 6.91 | 7.77 | 9.59 | 11.3 | 11.6 | 14.1 | 14.9 | 17.8 |
| 8 | 6.72 | 7.57 | 9.40 | 11.2 | 11.6 | 14.0 | 14.6 | 17.5 |
| 9 | 6.57 | 7.41 | 9.27 | 11.0 | 11.5 | 13.9 | 14.4 | 17.4 |
| 10 | 6.46 | 7.28 | 9.16 | 10.9 | 11.4 | 13.9 | 14.3 | 17.2 |
| **Average** | 7.64 | 8.57 | 10.3 | 12.1 | 12.2 | 14.7 | 15.8 | 18.8 |
| **Std-dev** | $\pm 1.3$ | $\pm 1.4$ | $\pm 1.3$ | $\pm 1.3$ | $\pm 1.1$ | $\pm 1.1$ | $\pm 1.6$ | $\pm 1.7$ |

**Table S15:** Diffusion Data (×10^−4^Å^2^*/*ps) for FF-wm5 using inear regression fitting over the MSD data (from 2-20 ps averaged over every 2 ps time-window)

|  |  |  |  |  |  |  |  |  |
| --- | --- | --- | --- | --- | --- | --- | --- | --- |
| **Aβ42** | 300 K | 310 K | 300 K | 310 K | 300 K | 310 K | 300 K | 310 K |
| **Shell** | **1–3 Å** |  | **3–4 Å** |  | **4–5 Å** |  | **5–8 Å** |  |
| 2 | 13.7 | 15.7 | 17.2 | 20.1 | 19.1 | 22.1 | 25.0 | 28.5 |
| 3 | 11.6 | 13.6 | 15.3 | 18.1 | 17.3 | 20.3 | 22.5 | 26.0 |
| 4 | 10.5 | 12.6 | 14.3 | 17.2 | 16.5 | 19.8 | 21.3 | 24.8 |
| 5 | 9.78 | 12.0 | 13.7 | 16.6 | 16.1 | 19.5 | 20.5 | 24.1 |
| 6 | 9.32 | 11.6 | 13.2 | 16.3 | 15.7 | 19.2 | 20.0 | 23.6 |
| 7 | 8.98 | 11.2 | 12.9 | 15.9 | 15.5 | 19.0 | 19.6 | 23.3 |
| 8 | 8.75 | 11.0 | 12.7 | 15.8 | 15.3 | 18.9 | 19.4 | 23.0 |
| 9 | 8.55 | 10.8 | 12.4 | 15.6 | 15.1 | 18.8 | 19.2 | 22.7 |
| 10 | 8.39 | 10.7 | 12.2 | 15.4 | 15.0 | 18.6 | 18.9 | 22.5 |
| **Average** | 9.95 | 12.1 | 13.8 | 16.8 | 16.2 | 19.6 | 20.7 | 24.3 |
| **Std-dev** | $\pm 1.7$ | $\pm 1.6$ | $\pm 1.6$ | $\pm 1.4$ | $\pm 1.3$ | $\pm 1.0$ | $\pm 1.9$ | $\pm 1.9$ |

| **Amylin** | 300 K | 310 K | 300 K | 310 K | 300 K | 310 K | 300 K | 310 K |
| --- | --- | --- | --- | --- | --- | --- | --- | --- |
| **Shell** | **1–3 Å** |  | **3–4 Å** |  | **4–5 Å** |  | **5–8 Å** |  |
| 2 | 13.9 | 15.8 | 17.2 | 20.9 | 19.1 | 22.7 | 24.7 | 28.7 |
| 3 | 11.7 | 13.6 | 15.3 | 18.8 | 17.3 | 20.8 | 22.2 | 26.1 |
| 4 | 10.6 | 12.5 | 14.3 | 17.8 | 16.5 | 20.1 | 20.8 | 24.7 |
| 5 | 9.94 | 11.8 | 13.7 | 17.0 | 16.1 | 19.7 | 20.0 | 24.0 |
| 6 | 9.44 | 11.4 | 13.4 | 16.6 | 15.7 | 19.3 | 19.5 | 23.5 |
| 7 | 9.10 | 11.0 | 13.0 | 16.2 | 15.6 | 19.3 | 19.0 | 23.2 |
| 8 | 8.82 | 10.8 | 12.7 | 16.1 | 15.2 | 18.9 | 18.6 | 22.9 |
| 9 | 8.62 | 10.5 | 12.5 | 15.8 | 15.1 | 18.3 | 18.4 | 22.6 |
| 10 | 8.44 | 10.4 | 12.4 | 15.6 | 15.0 | 18.3 | 18.2 | 22.3 |
| **Average** | 10.1 | 12.0 | 13.8 | 17.2 | 16.2 | 19.7 | 20.2 | 24.2 |
| **Std-dev** | ± 1.7 | ± 1.7 | ± 1.5 | ± 1.7 | ± 1.3 | ± 1.3 | ± 2.1 | ± 2.0 |
| **tau43** | 300 K | 310 K | 300 K | 310 K | 300 K | 310 K | 300 K | 310 K |
| **Shell** | **1–3 Å** |  | **3–4 Å** |  | **4–5 Å** |  | **5–8 Å** |  |
| 2 | 14.7 | 15.9 | 18.0 | 20.9 | 20.1 | 23.5 | 25.3 | 29.6 |
| 3 | 12.6 | 13.7 | 16.1 | 18.9 | 18.4 | 21.7 | 22.8 | 27.0 |
| 4 | 11.6 | 12.6 | 15.2 | 18.1 | 17.7 | 21.0 | 21.6 | 25.8 |
| 5 | 11.0 | 11.9 | 14.6 | 17.5 | 17.5 | 20.6 | 20.8 | 25.0 |
| 6 | 10.5 | 11.5 | 14.3 | 17.1 | 17.2 | 20.3 | 20.3 | 24.6 |
| 7 | 10.2 | 11.1 | 14.0 | 16.9 | 17.0 | 20.3 | 19.9 | 24.2 |
| 8 | 9.89 | 10.8 | 13.8 | 16.6 | 16.9 | 20.2 | 19.6 | 23.9 |
| 9 | 9.69 | 10.6 | 13.6 | 16.5 | 16.7 | 20.0 | 19.3 | 23.6 |
| 10 | 9.56 | 10.4 | 13.4 | 16.3 | 16.7 | 19.8 | 19.1 | 23.4 |
| **Average** | 11.1 | 12.1 | 14.8 | 17.6 | 17.6 | 20.8 | 21.0 | 25.2 |
| **Std-dev** | ± 1.7 | ± 1.7 | ± 1.4 | ± 1.4 | ± 1.0 | ± 1.1 | ± 1.9 | ± 1.9 |
| **αS** | 300 K | 310 K | 300 K | 310 K | 300 K | 310 K | 300 K | 310 K |
| **Shell** | **1–3 Å** |  | **3–4 Å** |  | **4–5 Å** |  | **5–8 Å** |  |
| 2 | 12.4 | 13.9 | 16.4 | 19.2 | 18.7 | 22.3 | 24.8 | 29.0 |
| 3 | 10.4 | 11.7 | 14.4 | 17.1 | 16.9 | 20.6 | 22.3 | 26.5 |
| 4 | 9.36 | 10.7 | 13.4 | 16.2 | 16.1 | 19.9 | 21.0 | 25.3 |
| 5 | 8.78 | 10.1 | 12.9 | 15.7 | 15.7 | 19.6 | 20.3 | 24.6 |
| 6 | 8.36 | 9.66 | 12.5 | 15.3 | 15.4 | 19.3 | 19.8 | 24.1 |
| 7 | 8.05 | 9.38 | 12.3 | 15.1 | 15.3 | 19.1 | 19.4 | 23.8 |
| 8 | 7.84 | 9.16 | 12.1 | 14.9 | 15.0 | 19.0 | 19.1 | 23.5 |
| 9 | 7.66 | 9.00 | 11.9 | 14.8 | 14.9 | 18.8 | 18.9 | 23.3 |
| 10 | 7.53 | 8.87 | 11.8 | 14.6 | 14.9 | 18.6 | 18.7 | 23.1 |
| **Average** | 8.93 | 10.3 | 13.1 | 15.9 | 15.9 | 19.7 | 20.5 | 24.8 |
| **Std-dev** | $\pm 1.5$ | $\pm 1.6$ | $\pm 1.4$ | $\pm 1.4$ | $\pm 1.2$ | $\pm 1.1$ | $\pm 1.9$ | $\pm 1.9$ |

**Table S16:** Diffusion Data (×10^−4^Å^2^*/*ps) for FF-wm3 using linear regression fitting over the MSD data (from 2-20 ps averaged over every 2 ps time-window)

|  |  |  |  |  |  |  |  |  |
| --- | --- | --- | --- | --- | --- | --- | --- | --- |
| **Aβ42** | 300 K | 310 K | 300 K | 310 K | 300 K | 310 K | 300 K | 310 K |
| **Shell** | **1–3 $\text{\AA}$** |  | **3–4 $\text{\AA}$** |  | **4–5 $\text{\AA}$** |  | **5–8 $\text{\AA}$** |  |
| 2 | 12.5 | 15.3 | 16.6 | 19.5 | 18.0 | 21.4 | 23.9 | 28.3 |
| 3 | 10.4 | 13.2 | 14.6 | 17.3 | 16.2 | 19.6 | 21.3 | 25.7 |
| 4 | 9.39 | 12.2 | 13.5 | 16.3 | 15.4 | 18.8 | 20.0 | 24.4 |
| 5 | 8.75 | 11.5 | 12.9 | 15.8 | 15.1 | 18.4 | 19.1 | 23.7 |
| 6 | 8.33 | 11.1 | 12.5 | 15.4 | 14.9 | 18.2 | 18.6 | 23.2 |
| 7 | 7.99 | 10.8 | 12.1 | 15.1 | 14.7 | 18.0 | 18.1 | 22.8 |
| 8 | 7.72 | 10.5 | 11.9 | 14.8 | 14.6 | 17.8 | 17.8 | 22.5 |
| 9 | 7.49 | 10.3 | 11.5 | 14.7 | 14.4 | 17.7 | 17.6 | 22.3 |
| 10 | 7.30 | 10.2 | 11.3 | 14.7 | 14.4 | 17.7 | 17.4 | 22.1 |
| **Average** | 8.87 | 11.7 | 13.0 | 15.9 | 15.3 | 18.6 | 19.3 | 23.9 |
| **Std-dev** | $\pm 1.6$ | $\pm 1.6$ | $\pm 1.7$ | $\pm 1.5$ | $\pm 1.1$ | $\pm 1.2$ | $\pm 2.1$ | $\pm 2.0$ |
| **tau43** | 300 K | 310 K | 300 K | 310 K | 300 K | 310 K | 300 K | 310 K |
| **Shell** | **1–3 $\text{\AA}$** |  | **3–4 $\text{\AA}$** |  | **4–5 $\text{\AA}$** |  | **5–8 $\text{\AA}$** |  |
| 2 | 13.6 | 16.0 | 17.6 | 20.5 | 19.6 | 22.7 | 25.3 | 28.9 |
| 3 | 11.4 | 13.9 | 15.6 | 18.6 | 18.0 | 21.0 | 22.8 | 26.4 |
| 4 | 10.4 | 12.8 | 14.8 | 17.6 | 17.4 | 20.4 | 21.5 | 25.1 |
| 5 | 9.86 | 12.1 | 14.3 | 17.1 | 17.0 | 20.0 | 20.7 | 24.4 |
| 6 | 9.47 | 11.7 | 13.9 | 16.7 | 16.7 | 19.6 | 20.2 | 23.9 |
| 7 | 9.15 | 11.3 | 13.7 | 16.5 | 16.5 | 19.4 | 19.8 | 23.5 |
| 8 | 8.89 | 11.0 | 13.5 | 16.3 | 16.5 | 19.3 | 19.5 | 23.2 |
| 9 | 8.68 | 10.8 | 13.3 | 16.1 | 16.3 | 19.4 | 19.3 | 22.9 |
| 10 | 8.51 | 10.6 | 13.0 | 16.0 | 16.4 | 19.2 | 19.1 | 22.6 |
| **Average** | 10.0 | 12.2 | 14.4 | 17.3 | 17.2 | 20.1 | 20.9 | 24.5 |
| **Std-dev** | $\pm 1.6$ | $\pm 1.7$ | $\pm 1.4$ | $\pm 1.4$ | $\pm 1.0$ | $\pm 1.1$ | $\pm 2.0$ | $\pm 2.0$ |
| **αS** | 300 K | 310 K | 300 K | 310 K | 300 K | 310 K | 300 K | 310 K |
| **Shell** | **1–3 $\text{\AA}$** |  | **3–4 $\text{\AA}$** |  | **4–5 $\text{\AA}$** |  | **5–8 $\text{\AA}$** |  |
| 2 | 12.1 | 13.7 | 16.4 | 19.2 | 18.8 | 22.2 | 24.9 | 29.2 |
| 3 | 10.1 | 11.5 | 14.4 | 17.2 | 17.0 | 20.5 | 22.4 | 26.5 |
| 4 | 9.10 | 10.4 | 13.5 | 16.3 | 16.3 | 19.8 | 21.2 | 25.2 |
| 5 | 8.50 | 9.72 | 13.0 | 15.7 | 15.9 | 19.4 | 20.5 | 24.5 |
| 6 | 8.12 | 9.29 | 12.6 | 15.3 | 15.6 | 19.2 | 20.0 | 24.0 |
| 7 | 7.84 | 8.97 | 12.3 | 15.0 | 15.5 | 18.9 | 19.7 | 23.6 |
| 8 | 7.59 | 8.72 | 12.1 | 14.8 | 15.2 | 18.7 | 19.4 | 23.4 |
| 9 | 7.39 | 8.51 | 12.0 | 14.6 | 15.2 | 18.6 | 19.1 | 23.1 |
| 10 | 7.22 | 8.37 | 11.9 | 14.5 | 15.1 | 18.6 | 19.0 | 23.0 |
| **Average** | 8.66 | 9.90 | 13.1 | 15.8 | 16.1 | 19.6 | 20.7 | 24.7 |
| **Std-dev** | $\pm 1.5$ | $\pm 1.7$ | $\pm 1.4$ | $\pm 1.5$ | $\pm 1.1$ | $\pm 1.1$ | $\pm 1.9$ | $\pm 2.0$ |

